# Structural diversity of the SARS-CoV-2 Omicron spike

**DOI:** 10.1101/2022.01.25.477784

**Authors:** Sophie M-C. Gobeil, Rory Henderson, Victoria Stalls, Katarzyna Janowska, Xiao Huang, Aaron May, Micah Speakman, Esther Beaudoin, Kartik Manne, Dapeng Li, Rob Parks, Maggie Barr, Margaret Deyton, Mitchell Martin, Katayoun Mansouri, Robert J. Edwards, Gregory D. Sempowski, Kevin O. Saunders, Kevin Wiehe, Wilton Williams, Bette Korber, Barton F. Haynes, Priyamvada Acharya

**Affiliations:** Duke Human Vaccine Institute, Durham NC 27710, USA; Duke University, Department of Medicine, Durham NC 27710, USA; Duke University, Department of Surgery, Durham NC 27710, USA; Theoretical Biology and Biophysics, Los Alamos National Laboratory, Los Alamos, NM 87545, USA; Duke University, Department of Immunology, Durham NC 27710, USA; Duke University, Department of Biochemistry, Durham NC 27710, USA

## Abstract

Aided by extensive spike protein mutation, the SARS-CoV-2 Omicron variant overtook the previously dominant Delta variant. Spike conformation plays an essential role in SARS-CoV-2 evolution via changes in receptor binding domain (RBD) and neutralizing antibody epitope presentation affecting virus transmissibility and immune evasion. Here, we determine cryo-EM structures of the Omicron and Delta spikes to understand the conformational impacts of mutations in each. The Omicron spike structure revealed an unusually tightly packed RBD organization with long range impacts that were not observed in the Delta spike. Binding and crystallography revealed increased flexibility at the functionally critical fusion peptide site in the Omicron spike. These results reveal a highly evolved Omicron spike architecture with possible impacts on its high levels of immune evasion and transmissibility.

## Introduction

The SARS-CoV-2 Omicron (B.1.1.529) variant was identified November 24^th^, 2021 in South Africa, declared a Variant of Concern (VOC) by the World Health Organization on November 26^th^, and has rapidly replaced Delta (B.1.617.2) as the dominant form of SARS-CoV-2 circulating globally. The Omicron spike (S) protein harbors 30 mutations and is the most immune evasive VOC identified thus far, surpassing Beta (B.1.351) in its ability to resist neutralization by antibodies (Abs) (Supplementary Figure 1) (*1–4*). Structural studies have been instrumental in revealing changes in VOC S protein conformations, and in understanding the atomic level mechanisms that drive higher transmissibility and immune evasion (*5–9*).

**Figure 1.**
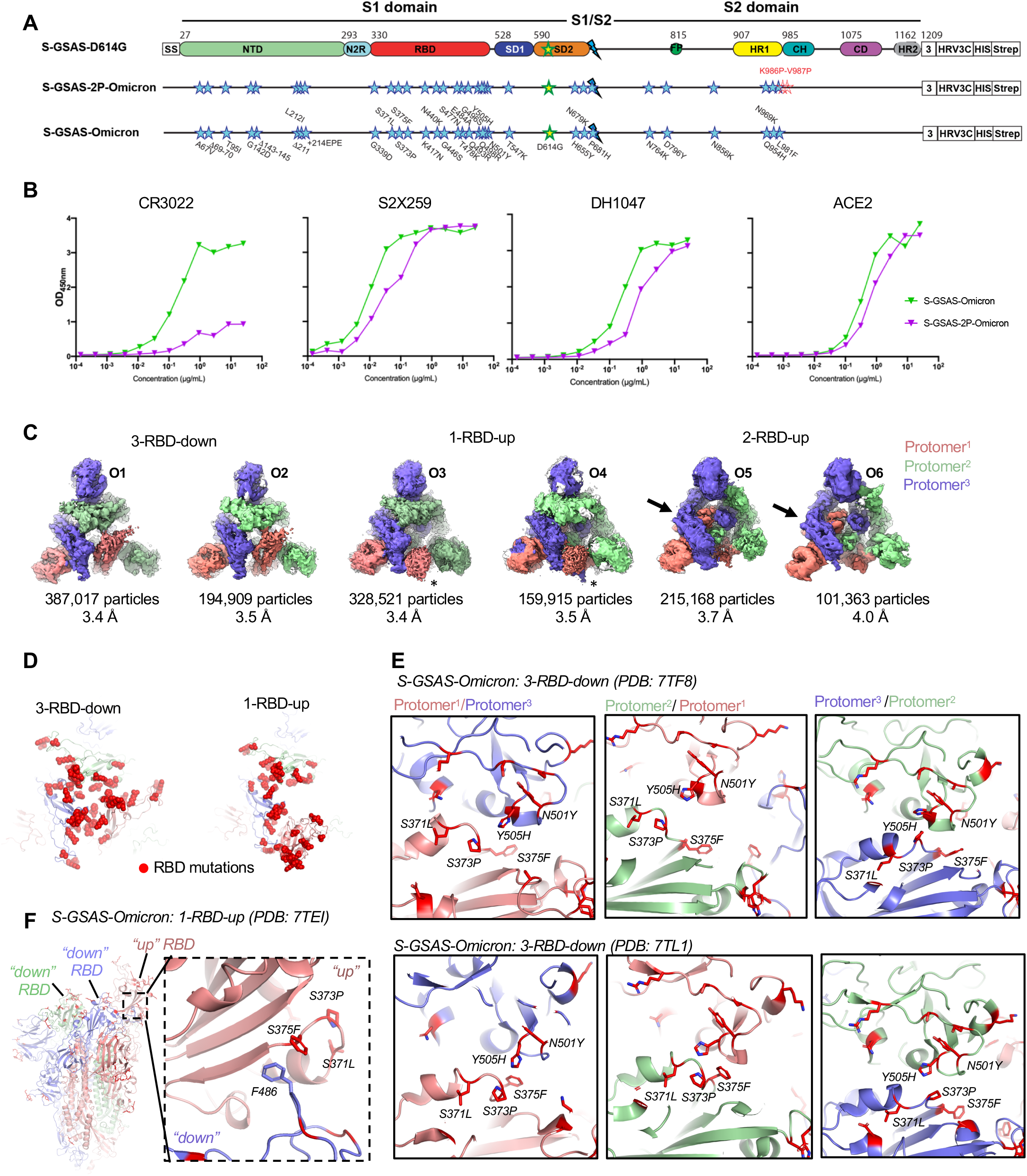
Conformational Diversity of the SARS-CoV-2 Omicron (B.1.1.529) S protein. **(A)** Domain architecture of the SARS-CoV-2 S protomer. The S1 subunit contains a signal sequence (SS), the NTD (N-terminal domain, pale green), N2R (NTD-to-RBD linker, cyan), RBD (receptor-binding domain, red), and SD1 and SD2 (subdomains 1 and 2, dark blue and orange) subdomains. The S2 subunit contains the FP (fusion peptide, dark green), HR1 (heptad repeat 1, yellow), CH (central helix, teal), CD (connector domain, purple), and HR2 (heptad repeat 2, gray) subdomains. The transmembrane domain (TM) and cytoplasmic tail (CT) are replaced by a foldon trimerization sequence, followed by a HRV3C cleavage site (HRV3C), a his-tag (His), and a strep-tag (Strep). The D614G mutation in the SD2 domain is indicated by a yellow star with green contour. The location of the S1/S2 furin cleavage site (RRAR mutated to GSAS) is indicated by a blue lightning sign. The K986P-V987P mutations between the HR1 and CH domains in S-GSAS-2P-Omicron are indicated by red stars. **(B)** Binding of S-GSAS-Omicron and S-GSAS-2P-Omicron to RBD-directed antibodies CR3022, S2X259 and DH1047, and to ACE2 receptor. **(C)** Cryo-EM reconstructions of Omicron S protein 3-RBD-down (O1: EMD-25865, PDB 7TF8; O2: EMD-25983, PDB 7TL1), 1-RBD-up (O3: EMD- 25846, PDB 7TEI; O4: EMD-25984, PDB 7TL9) and 2-RBD-up (O5: EMD-25880, PDB 7TGE; O4) states, colored by protomer, and viewed down from the host cell membrane. In the 1-RBD-up reconstructions, the “up” RBD is indicated by an asterisk (*). The black arrows in the 2-RBD-up protomer point to the “down” RBD. **(D)** Omicron spike 3-RBD-down (O1: EMD-25865, PDB 7TF8) and 1-RBD-up (O3: EMD-25846, PDB 7TEI) structures, colored by promoter, with RBD mutations shown in red spheres. **(E and F)** Interprotomer RBD-RBD interactions in **(E)** the 3-RBD-down state, and **(F)** the 1-RBD-up state.

The pre-fusion SARS-CoV-2 S protein is composed of the S1 and S2 subunits that undergo structural changes to facilitate receptor binding and fusion with the host-cell membrane (*10, 11*). While the S2 subunit is conformationally stable prior to receptor engagement, the S1 subunit, with its mobile N-terminal domain (NTD) and receptor-binding domain (RBD), is inherently dynamic. The RBD transitions between a “closed” (“down”) state where the binding site for the ACE2 receptor is occluded, and an “open” (“up”) state that exposes the ACE2 biding site (*10, 11*). Following receptor binding and proteolytic processing of the S protein, the S2 subunit undergoes large conformational changes that result in release of the fusion peptide (FP) to mediate fusion of the virus and host-cell membranes. RBD dynamics are impacted by interprotomer RBD-to-RBD and RBD-to-NTD contacts, as well as by other S protein structural units, including the SD1 and SD2 subdomains, and the “N2R (NTD-to-RBD) linker” that connects the NTD and RBD within a protomer. We previously described how the S1 domain interactions are modulated by VOCs to alter the S protein RBD presentation and how these can be exploited for immunogen design (*5, 12, 13*).

Here, we determine structures of native, unstabilized Omicron and Delta S protein ectodomains to understand how the acquired mutations alter their conformational states, and influence receptor binding site and Ab epitope presentation. The S ectodomains were prepared in our previously described S-GSAS-D614G platform without S2 subunit proline stabilization mutations as these can alter the conformational landscape of the pre-fusion S protein (Fig. 1A, fig. S2) (*12, 14*). We determined structures of 3-RBD-down, 1-RBD-up and 2-RBD-up populations of the Omicron and Delta S ectodomains (Fig. 1–3, figs. S3 to S7). We observed considerable down-state variability in the Delta S1 subunit with one structural class presenting a disordered S1 subunit protomer, similar to an ectodomain structure we previously described in a mink-associated spike (*5*). In contrast, the Omicron S protein displayed reduced S1 variability with several of its sixteen immune-evasive RBD amino acid substitutions stabilizing the RBD-RBD interfaces. These substitutions also stabilized 1-RBD-up states in a manner not observed in the Delta spike. The tight packing of the RBDs in the Omicron 3-RBD-down structures limits RBD motion, causing a prominent single protomer NTD-to-RBD (N2R) linker rearranged conformation. This rearrangement was identified in Delta and other variants but was comparatively rare among them. Altered S2 conformational dynamics was indicated by weaker binding of Omicron S protein (relative to the G614 and Delta S proteins) to 2G12 and other Fab-dimerized glycan-reactive (FDG) Abs that target a quaternary S2 glycan cluster, as well as by enhanced binding to fusion peptide (FP) directed Ab DH1058. A high-resolution crystal structure of the FP-Ab complex suggests enhancement must occur through release of the FP from its closed state position. Enhanced FP dynamics may therefore be linked to Omicron’s enhanced transmissibility. Together, these results point to an Omicron spike evolved beyond immune evasion toward a more compact architecture with a well-regulated fusion machinery and altered dynamics of its fusion peptide (FP) that leads to more facile FP release.

## Results

### Conformations of the SARS-CoV-2 Omicron S protein

As recent studies have reported structures of the Omicron S protein with Proline stabilizing mutations in the S2 subunit (*15–18*), either the 2P or the HexaPro mutations (*10, 14*), we first assessed the effect of such stabilization by comparing binding of the S-GSAS-Omicron and S-GSAS-2P-Omicron to the receptor ACE2 and RBD-directed Abs (Fig. 1, A and B). We found that CR3022, an Ab targeting a cryptic RBD epitope (*19*) lost >90% binding to S-GSAS-2P-Omicron, suggesting that the 2P mutations are limiting the conformational diversity of the Omicron S protein ectodomain. Reduced binding for the 2P vs non-2P Omicron spike is also seen for the 3-RBD-up conformation binding DH1047 and S2X259 Abs (*20, 21*). We therefore determined structures of the native Omicron S protein ectodomain in the S-GSAS form to examine the impacts mutations had on its conformation (Fig. 1, fig. S2 to S5, table S1). In the cryo-EM dataset, we identified 3-RBD-down, 1-RBD-up and 2-RBD-up populations of the S protein ectodomain (Fig. 1C). The Omicron spike harbors 16 amino acid substitutions in the RBD (Fig.1D), of which several have been shown to mediate ACE2 recognition and/or immune escape (*3, 22*). The 3-RBD-down populations classified into two asymmetric reconstructions with each displaying close RBD-RBD pairing. Of the two 3-RBD-down structures, named O1 and O2, the interprotomer domain arrangement appeared more symmetric in O2 than in O1. This asymmetry was visualized in difference distance matrices (DDMs) that provide superposition-free comparisons between a pair of structures by calculating the differences between the distances of each pair of Ca atoms in a structure and the corresponding pair of Ca atoms in the second structure (fig. S5). We assigned the tag Protomer^1^ to the protomer with the most mobile RBD in each reconstruction, characterized by the weakest RBD map density (Fig. 1C). Interprotomer interactions between the “down” RBDs were mediated by a loop containing 3 amino acid substitutions, S371L, S373P and S375F. A rearrangement of the loop caused by the S373P mutation facilitated closer packing of the RBD-RBD interface via interactions of S373P and S375F with the N501Y and Y505H substitutions in the adjacent RBD (Fig. 1E). In the 1-RBD-up structure the S375F substitution created an interprotomer interaction with residue Phe486 of the adjacent RBD-down protomer (Fig. 1F). In the 2-RBD-up state, both up RBDs were disordered, indicating increased mobility (Fig. 1C). Overall, these results show that the mutations in the Omicron S protein induced coupling of the RBDs causing unique S protein closed and open structural states.

We previously showed that the RBD up/down transitions are accompanied by movements of the SD1/SD2 subdomains, and of the N2R linker (residues 306-334) that connects the NTD and RBD within a single protomer unit (Fig. 2A) (*12*). In the RBD “down” state the N2R region stacks against and contributes a β strand to each SD1 and SD2 subdomain (Fig. 2B). Notably, in the Omicron 3-RBD-down structure O1 (PDB ID: 7TF8) (Fig. 1C and 2B), we found that the N2R region secondary structures in the mobile Protomer^1^ was modified, with a break in the SD2-associated β strand stabilized by a new intraprotomer salt bridge formed between R319 and NTD residue E298 (Fig. 2, B and C). This was accompanied by packing of residue F318 against the NTD 295-303 helix. This rearrangement permits interaction with a typically disordered segment of SD2, named “SD2 flex” here (residues 619-642). Stabilization of SD2 flex was facilitated by Van der Waals interactions of residue Ile624 (SD2 flex) with SD2 residues V595 and Y612 (Figure 2C) and Trp633 packing between NTD residue P295 and N2R linker residue R319. The spatial positioning of the NTD helix 295-303, of the SD2 loop and of the SD1 loop 554-565 were similar between the “up” protomer (Protomer^1^) of the 1-RBD-up structure and the mobile protomer (Protomer^1^) of the 3-RBD-down structure, suggesting that the mobile protomer may be poised to transition to the “up” position. The mobile SD1 region was stabilized in both the 3-RBD-down and 1-RBD-up Omicron S proteins via interprotomer hydrogen bonding with the S2 subunit mediated by residues D568 and T572 of SD1 with the N856K amino acid substitution near the Omicron S protein fusion peptide (Fig. 2, E and F). A hydrogen bond also formed between the carboxyl oxygen of residue T315 of the N2R linker in the “up” protomer with the N764K substitution in the Omicron S protein. Thus, strategically placed residue substitutions in the Omicron S protein stabilize the highly mobile regions in the S1 subunit. Together, the close packing of the RBDs in the 3-RBD-down Omicron S protein and the N2R rearrangement in the “down” protomers define the wide range of conformational impacts that emanate from the extensive network of its acquired mutations.

**Figure 2.**
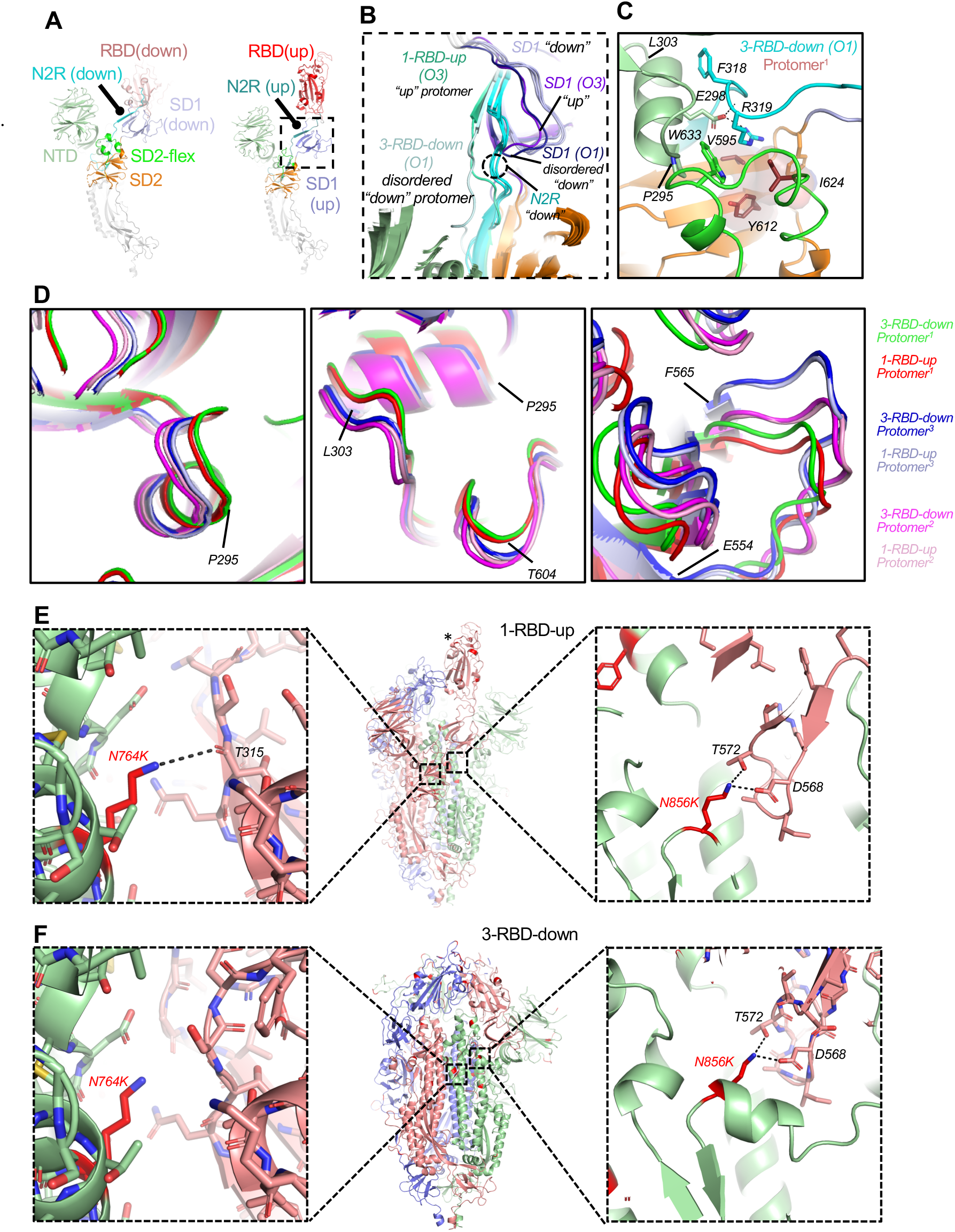
Intraprotomer communication through the NTD-to-RBD (N2R) linker. **(A)** Single protomers shown for RBD-down (left) and RBD-up (right) states, colored by domains: NTD -pale green, “up” RBD –red, “down” RBD – salmon, SD1 subdomain colored light and dark blue, respectively for the up and down protomer, N2R linker colored cyan for down protomer and deep teal for up protomer. **(B)** Zoomed in view of the N2R and surrounding regions showing overlay of all protomers. The N2R region is colored sea green for the RBD-up protomer (Protomer^1^) in the 1-RBD-up structure O3, light cyan for the disordered RBD protomer (Protomer^1^) in the 3-RBD-down structure O1, and cyan for all the “down” RBD protomers (Protomer^2^ and Protomer^3^). SD1 is colored pale blue for all “down” protomers in O1 and O3 (Protomer^2^ and Protomer^3^), purple and dark purple for Protomer^1^ of O3 and O1, respectively. **(C)** Zoomed-in view of the interaction of the N2R linker of Protomer^1^ of the 3-RBD-down state O1 with the NTD helix spanning residue 295-303. N2R linker is colored cyan, NTD pale green, SD2 orange, SD2 flex green. Brown surface shows interaction between V595, Y612, and I624. **(D)** Overlay of protomers showing views of the NTD helix 295-303 (left), SD2 loop centered on T604 (middle) and SD1 region 554-565. Protomers are colored as indicated. Superpositions were performed using S2 residues 908-1035. **(E)** Stabilization of the SD1 and N2R linker regions in the “up” protomer of the 1-RBD-up structure via hydrogen bonds acquired through residue substitutions in the Omicron S protein. **(F)** Stabilization of the SD1 region in the 3-RBD-down structure via hydrogen bonds acquired through residue substitutions in the Omicron S protein.

### Conformations of the SARS-CoV-2 Delta S protein

We next studied the Delta S protein to understand the differences in its structural properties that may underlie the differences in its pathobiology with Omicron. We determined cryo-EM structures of the S-GSAS-Delta S protein ectodomain (Fig. 3, figs. S6 and S7, and table S1). The Delta variant S protein includes two substitutions and a deletion in the NTD, two RBD substitutions, a P681R substitution proximal to the furin cleavage site, and a D950N substitution in the HR1 region of the S2 subunit (Fig. 3A and fig. S1). We identified 3-RBD-down (D1), 1-RBD-up (D2) and 2-RBD-up (D3) S protein ectodomain populations (Fig. 3A), as in the Omicron dataset, in addition to a population, named D4, which exhibited very high disorder in one of its S1 subunits such that the entire S1 subunit, including the NTD, RBD, SD1, N2R linker, as well as part of the SD2 subdomain, were not visible in the cryo-EM reconstruction (Fig. 3A). This was similar to a state found in a mink-associated spike (*5*). The 3-RBD-down population classified into seven distinct subclasses (D5-D10) (Fig. 3B), of which one (D6) displayed an N2R configuration observed in the Omicron O1 Protomer^1^ (Fig. 3C). A similar dislocation of the N2R region from its β-strand arrangement with the SD2 subdomain was also found in a “down” protomer of a 1-RBD-up subclass (D12) (Fig. 3C), possibly representing an intermediate to the 2-RBD-up state. The array of distinct populations of the Delta S ectodomain that differ in their S1 subunit conformation are reminiscent of our observations with other naturally-occurring variants (*5, 12*), while the appearance of the single S1-protomer disordered state (Fig. 3, A and D) as we had observed in a mink-associated spike hints of S protein instability originating in the mobile S1 region encompassing the NTD, N2R linker, RBD and SD1.

**Figure 3.**
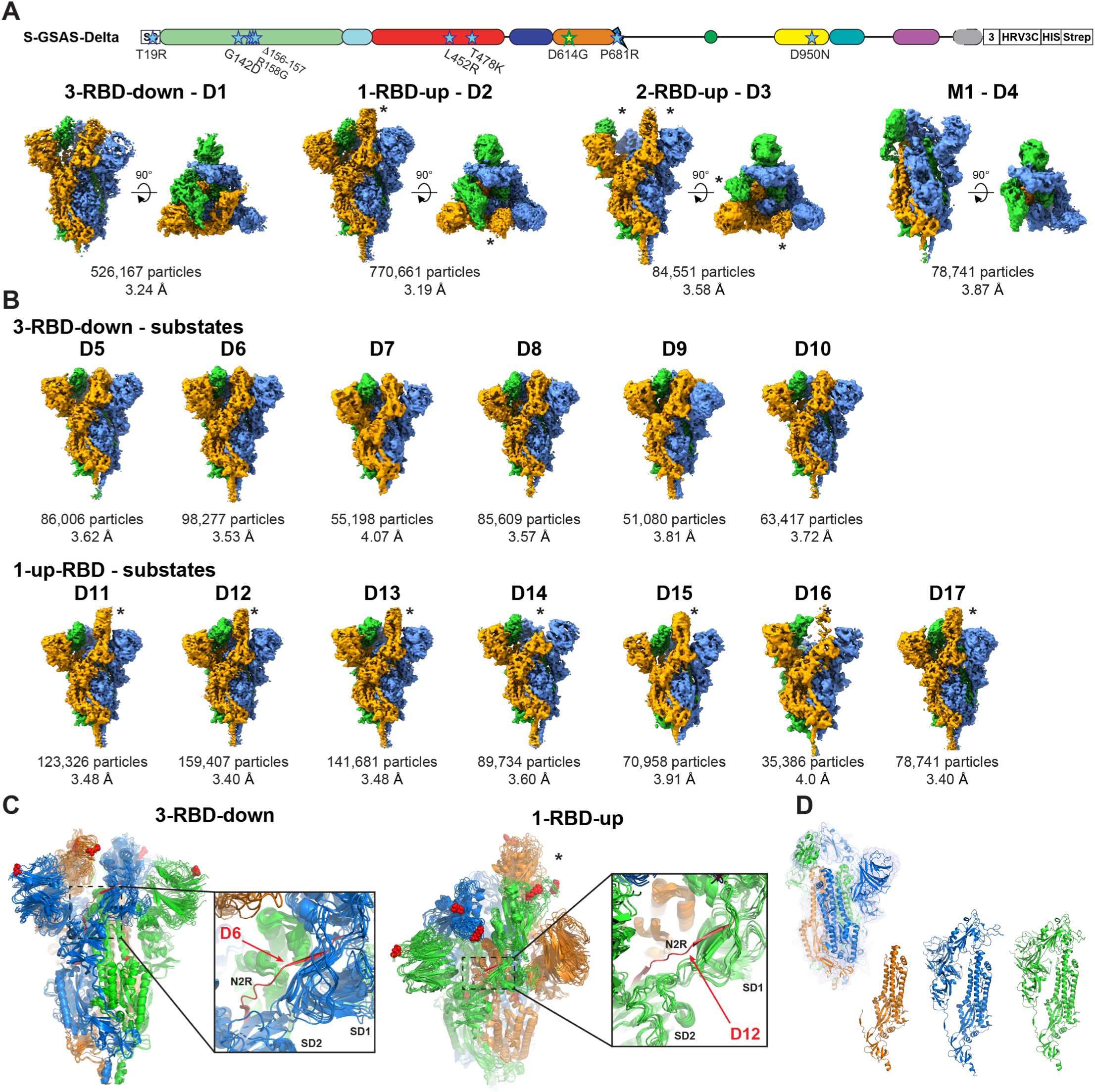
Conformational Diversity of the SARS-CoV-2 Delta (B.1.617.2) S protein. **(A)** Top. Domain architecture of the SARS-CoV-2 Delta S protomer. Bottom.Side views and top views of the Delta S protein cryo-EM reconstructions of 3-RBD-down, 1-RBD-up, 2-RBD-up and M1 states, colored by protomer. The “up” RBD are indicated by an asterisk (*). **(B)** Side views of subpopulations obtained by further classification of the states shown in **(A)**. **(C)** Left. Overlay of 3-RBD-down subclasses. Dashed rectangle indicates the N2R region zoomed in on the right image. The N2R linker in one of the Delta 3-RBD-down classes showed a N2R region that was dislodged from its beta strand arrangement with the SD2 subdomain, and is shown in red. Right. Overlay of 1-RBD-up subclasses. Dashed rectangle indicates the N2R region zoomed in on the right image. The N2R linker in one of the two “down” protomers of a 1-RBD-up structure showed a N2R region that was dislodged from its beta strand arrangement with the SD2 subdomain, and is shown in red. Mutations in the Delta variant are shown as red spheres **(D)** M1 state with the S1 subunit and SD2 subdomain of one of the three protomers disordered. The cryo-EM density is shown as a blue mesh with the underlying fitted model colored by protomer. The protomer colored orange shows disorder in the S1 subunit and SD2 subdomain, therefore, these regions could not be built in this protomer.

### Domain positioning in Omicron, Delta and engineered S proteins

We next investigated the source of the N2R linker rearrangements in the Delta and Omicron 3-RBD-down structures. The S1 subunit domain arrangements are responsive to one another across protomers through communication between adjacent, contacting RBDs, NTDs and subdomains. We have demonstrated through engineering (*13*) and by examination of previous variants (*5, 12*) that this communication plays an essential role in RBD up/down state presentation. The close interaction between RBDs in the Omicron spike led us to ask whether a design engineered to lock the 3-RBD-down state, termed rS2d for the introduced RBD to S2 disulfide staple, would show a similar N2R linker rearrangement due to restrictions on RBD movement. We obtained cryo-EM reconstructions of an S2 stabilized HexaPro version of the rS2d design (rS2d-HexaPro) and for the unstabilized rS2d (figs. S8 and S9, table S1) (*13, 14, 23*). Three-dimensional classification of both datasets led to two prominent structural states (Fig. 4). One reconstruction in both datasets, each referred to as State 1, displayed a similar SD2 rearrangement as that observed in the Delta D6 and the Omicron O1 3-RBD-down structures (Figure 4A). Examination of a structural morph between both states (Video S1) identified a marked shift in the SD1 toward SD2 in the linker displaced protomer (Fig. 4, B and C). Because of the proximity between the two subdomains, the β-sheet secondary structure linking the two is broken which in turn breaks the paired N2R β-sheet structure (Fig. 4B). The loss of this secondary structure permits the observed rearrangement in the linkers and the disordered segment.

**Figure 4.**
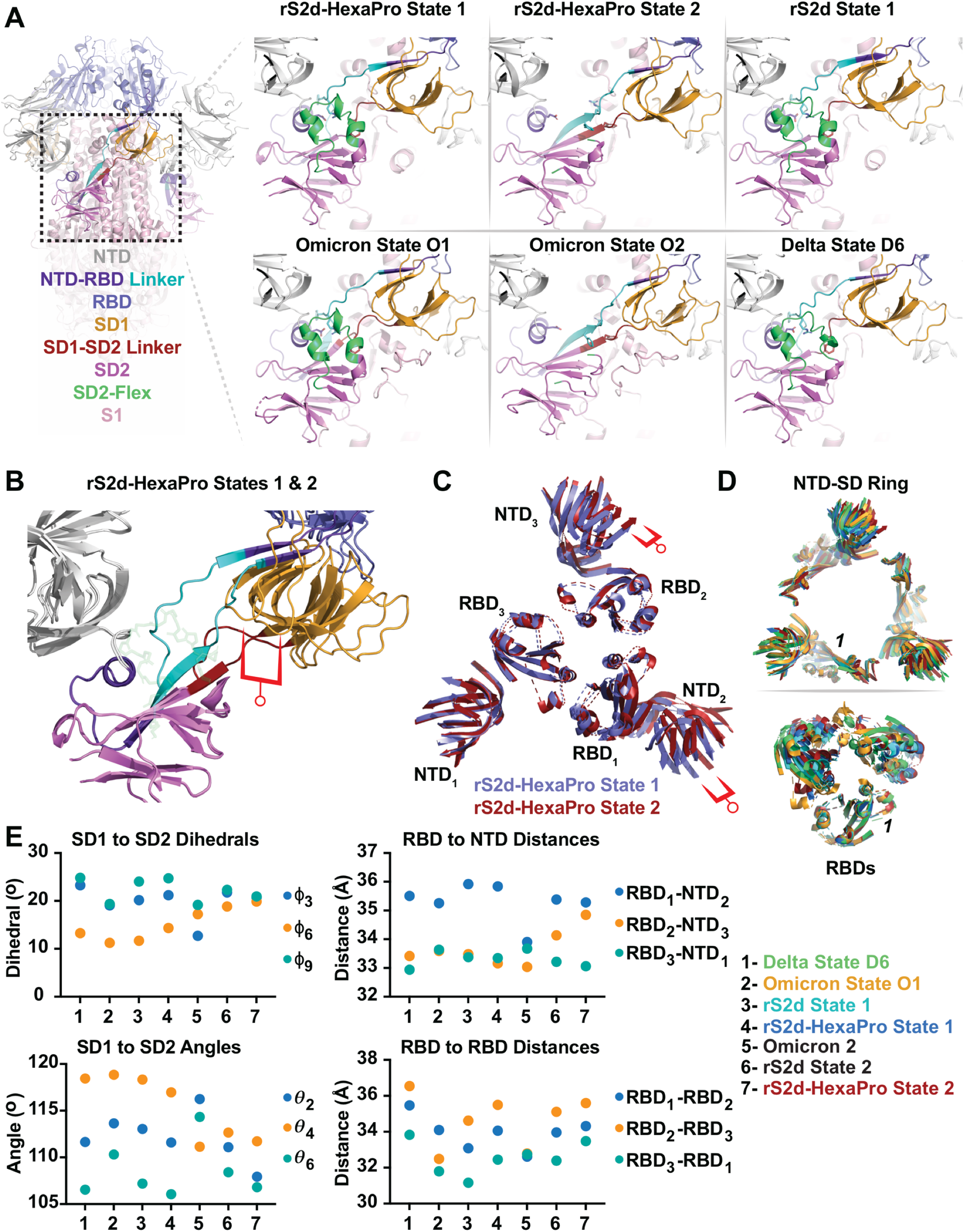
The down state locked rS2d trimer is structurally similar to the Omicron 3-RBD-down S protein. **(A)**(left) Spike trimer ectodomain highlighting the SD2 region. (right) Structural states of the rS2d, rS2d-HexaPro, Omicron, and Delta state D6 Spikes in the region of SD2 highlighting structural rearrangements in the linking regions (cyan and dark red) and flexible SD2 segment (green). **(B)** Comparison of the SD1 to SD2 disposition between rS2d-HexaPro states 1 and 2 via alignment of the SD2 core residues. The red indicator highlights the distance shift in SD1. **(C)** NTD aligned RBD pairings between rS2d-HexaPro states 1 and 2. The red indicator highlights the difference in RBD disposition relative to the NTD in the RBD2 to NTD3 pairing. Only sheet and helix elements are shown for clarity. **(D)** (top) Alignment of the NTDs and subdomains of the rS2d-HexaPro states 1 and 2, rS2d state 1, Omicron, and Delta D6 trimers. (bottom) Alignment of the rS2d-HexaPro states 1 and 2, rS2d state 1, Omicron, and Delta D6 trimer RBDs. Only sheet and helix elements are shown for clarity. Number indicates protomer 1 position. **(E)** Vector dihedral, angle, and distances defining differences and similarities between the rS2d-HexaPro states 1 and 2, rS2d states 1 and 2, Omicron, and Delta D6 trimers. Phi angles 3, 6, and 9 and Theta angles 2, 4, and 6 correspond to dihedrals/angles in protomers 3, 1, and 2, respectively.

We next asked whether the features in the rS2d spike domain arrangements that led to N2R linker rearrangement occurred in the Delta and Omicron 3-RBD-down states. Alignment of the SD2 subdomains of the N2R rearranged protomers indicated the overall subdomain and NTD domain arrangements were similar, except for the State 2 protomers (Figure 4D). Alignment of the disordered RBD (Protomer^1^) revealed the RBD positions differed markedly among the trimers (Fig. 4D). Using a vector-based quantification of S protein domain arrangement (*5, 13*), we previously found that absolute positions in spike domains may differ, but that trends in their overall architecture are correlated with important structural features such as the propensity to occupy the RBD-up state. The most important feature observed in the SD2-rearranged rS2d state was forcing of the SD1 toward its adjacent SD2.

We examined a series of vectors connecting the subdomains and NTDs of each protomer (Fig. 5A) finding that, while the angle and distance values differ slightly, the trend in relative positions are the same for each of the S protein trimers where the N2R rearrangement was observed (Fig. 4E). Specifically, the shift in SD1 position toward SD2 relative to the rS2d and rS2d-HexaPro State 2 is retained, consistent with a role of these subdomain shifts in causing the N2R rearranged state. These local effects were accompanied by a reduced distance between the RBD-NTD pair across from the disordered RBD protomer (RBD_2_ to NTD_3_) suggesting, despite variability in RBD positions, this interaction plays a role in the N2R rearrangement (Fig. 4E). Though the N2R rearranged trimers were similar, the non-rearranged State 2 trimers of the Omicron and rS2d trimers differed markedly. Together, comparison of the rS2d constructs and the variant structures indicates local and global rearrangements in S1 lead to the rearranged state.

**Figure 5.**
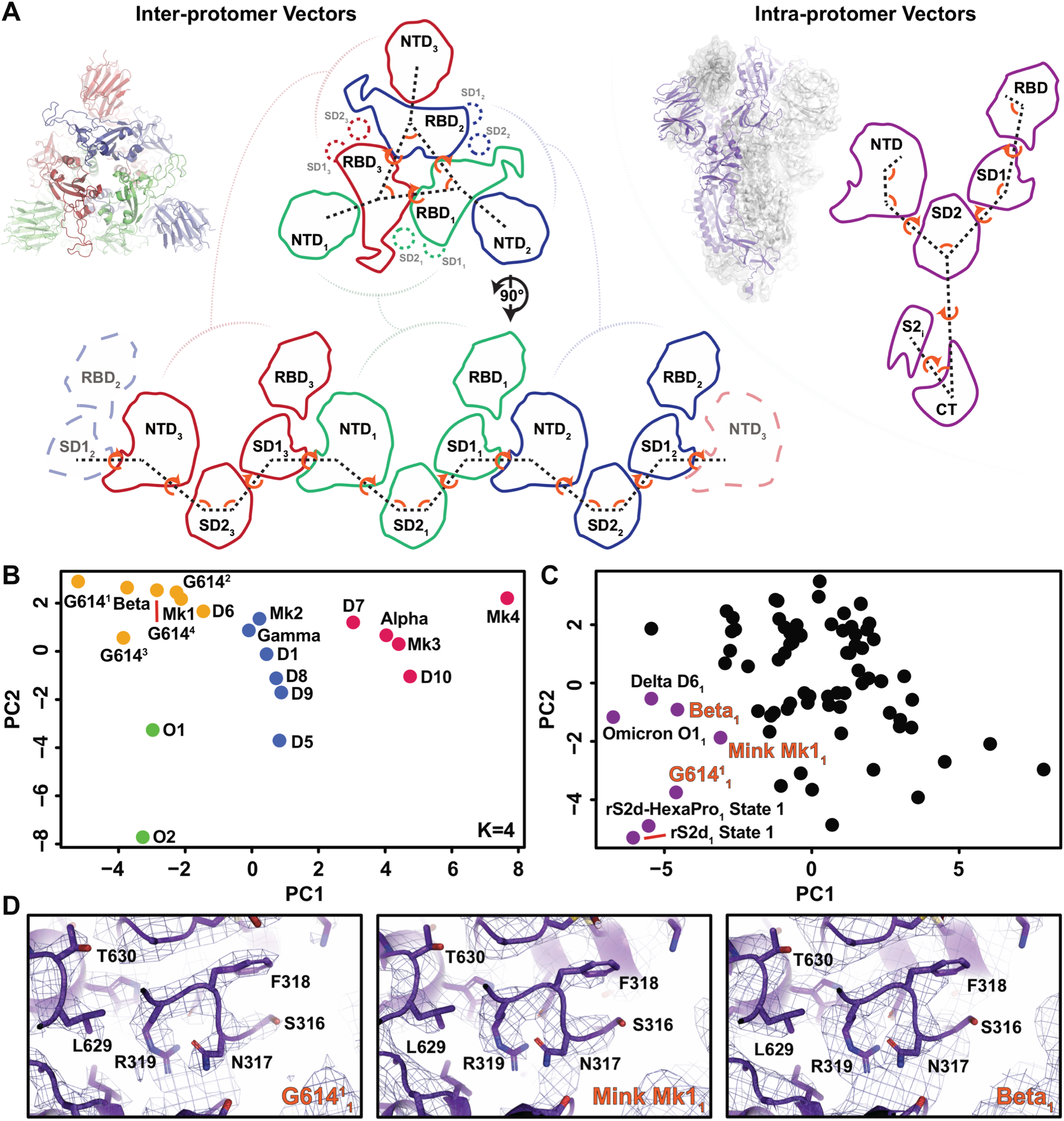
Intra- and interprotomer domain relationships in SARS-CoV-2 3-RBD-down S proteins. **(A)** (left) Inter-protomer vectors describing the relationship between the NTDs, RBDs, and subdomains across protomers. (right) Intra-protomer vectors describing the within protomer domain geometries. **(B)** Principal components analysis of the inter-protomer vectors for each variant structure. Colors indicate K-means centers using a total of four centers. The M notation indicates values from previously determined structures of the cluster-5, Mink associated Spike (PDB IDs 7LWL, 7LWI, 7LWK, 7LWJ for M1-4, respectively) while the G164 superscript notation indicates values from previously determined structures for the D614G Spike (PDB IDs 7KE8, 7KE6, 7KE7, and 7KE4 for G6141-4, respectively). **(C)**Principal components analysis of the intra-protomer vectors for each variant structure protomer. Protomers displaying the N2R rearrangement are highlighted in purple. **(D)** Previously determined cryo-EM maps of the principal components analysis identified N2R rearranged protomers aligned with the State 1 rS2d N2R rearranged protomer.

### Vector analysis of intra- and interprotomer domain relationships in the Omicron and Delta S proteins

We next examined clustering of the Omicron and Delta variant 3-RBD-down structures with previous variant structures utilizing sets of interprotomer and intraprotomer vectors using principal components analysis (PCA) (Fig. 5A). As suggested by visual inspection of the Omicron variant structures, the Omicron trimers form a distinct cluster in PCA of the interprotomer vector set (Fig. 5B). The Delta structures were spread throughout the variant structure set along principal component one, consistent with the considerable structural variability recovered through sub classification of the cryo-EM dataset (Figure 3). These results demonstrate the considerable structural rearrangements that the S protein has acquired through SARS-CoV-2 evolution (Fig. 5B). The structural conservation of the rearranged N2R state between the engineered and variant S proteins suggests this plays a specific role in S1 dynamics. As a specific domain organization accompanies the appearance of this state, we asked whether previous variant structures may present this state, albeit at lower proportions of the total population and with less conspicuous map density. We therefore examined the intraprotomer vector set by PCA to identify candidate structures (Fig. 5, A and C). Consistent with the interprotomer vectors, N2R rearranged protomers of the rS2d constructs and the Omicron and Delta variants occupied a distinct position along principal component one (Fig. 5C). Protomers from the D614G, Mink, and Beta variants clustered with these structures. Densities in the N2R region for these structures were consistent with the rearranged state as determined by fitting of the rS2d State 1 coordinates, where this N2R rearranged state was particularly well-resolved, to the respective cryo-EM densities (Fig. 5D). As expected, the densities were less clear than in the rS2d constructs, suggestive of multistate or dynamic behavior. As was observed in the rS2d, Omicron and Delta structures, the rearranged N2R protomer contained the disordered RBD and corresponded to the most distant RBD-NTD pair. These results highlight the differences in the Omicron 3-RBD-down structures from other variant S proteins, characterized by the stabilization of a N2R rearranged state.

### Antigenicity of the Omicron and Delta S proteins

We next studied the antigenicity and receptor binding properties of the Omicron S protein. Consistent with the extensive immune escape observed with the Omicron variant (*1–4, 22*), we found that its S protein lost binding to several SARS-CoV-2 neutralizing Abs (Fig. 6, A to C, fig. S10). Ab DH1050.1 that targets a site of vulnerability in the NTD (*20, 24*), no longer bound the Delta and Omicron S proteins (Fig. 6, A and B). The non-neutralizing, protective Ab DH1052 retained binding to the Delta but not to the Omicron S protein, its binding likely disrupted by changes in the region spanning residues 211-215 of the Omicron S protein (Fig. 6, A and B) (*20*). The RBD receptor binding site directed Abs DH1041 and DH1042 lost binding to Omicron S protein while DH1042 also lost binding to Delta S protein. Both Omicron and Delta S proteins retained similar binding levels to ACE2 (Fig. 6, A and C, fig. S10). DH1041 and DH1042 bind similar RBD epitopes overlapping the ACE2 binding site (Fig. 6C, figs. S11 and S12) (*20*), showing that subtle changes in epitope footprint can alter the susceptibility profile of Abs to residue substitutions. For DH1042, insertion of a charged residue within a hydrophobic binding site by the L452R substitution in the Delta S protein (fig. S12) results in reduction of binding and loss in neutralization activity. Indeed, DH1042 showed substantially reduced binding to S-GSAS-L452R and the S-GSAS-Epsilon S protein ectodomain (B.1.429) that harbors the L452R substitution. DH1041 binding, on the other hand, is unaffected by the L452R substitution (fig. S12).

**Figure 6.**
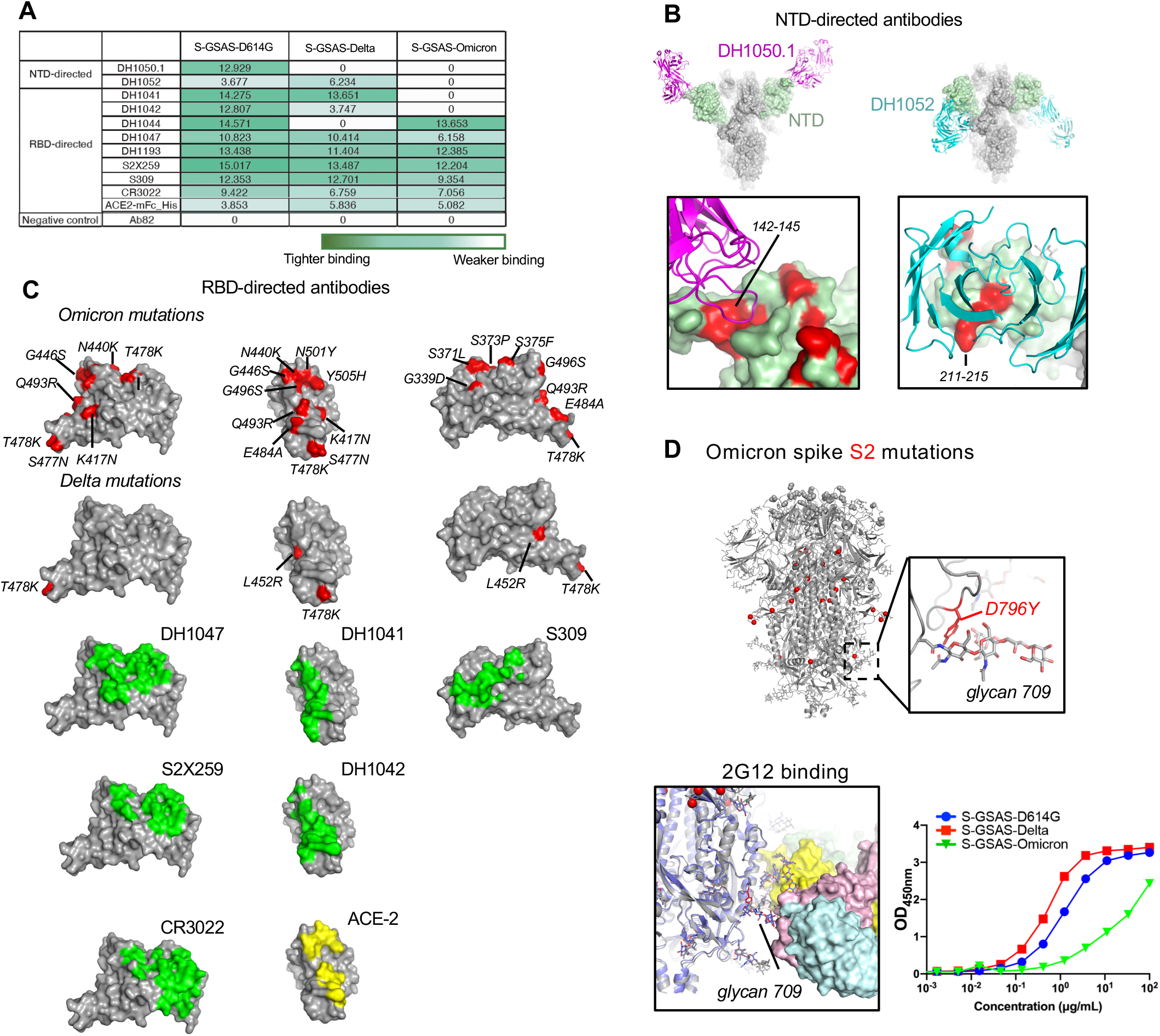
Impact of Delta and Omicron mutations on the antigenicity of the SARS-CoV-2 S protein. **(A)** Antibody binding to SARS-CoV-2 S proteins measured by ELISA. The binding values were obtained by calculated area under curve of ELISA binding curves. **(B)** Binding of NTD directed antibodies DH1050.1 and DH1052 to the SARS-CoV-2 spike. **(C)** Top two rows show three views of the RBD with residues mutated in Omicron (top row) and Delta (second row) colored red. Binding sites of RBD-directed antibodies and ACE2 receptor to SARS-CoV-2 S ectodomains are shown). **(D)**Top. Omicron 3-RBD-down spike is shown in grey, with residue changes relative to the D614G variant shown as spheres. Residue changes in the S2 region are colored red. The zoomed in image shows glycan 709 and its interaction with the D796Y substitution. Bottom. Binding of 2G12 to SARS-CoV-2 S proteins.

Consistent with its reported retention of Omicron neutralization, albeit at reduced levels, Ab S309, the parental form of the engineered therapeutic Ab Sotrovimab, retained substantial binding to the Omicron spike (Fig. 6A and fig. S10). The broad sarbecovirus neutralizing Ab DH1047 (*20, 25*) lost substantial binding to the Omicron spike, resulting in loss in its neutralization activity against Omicron (*26*). The S2X259 Ab targets a similar epitope (*21*) but retained binding and neutralization activity against Omicron (*3*). The SARS-CoV-2 cross-reactive Ab CR3022 targets a cryptic, unmutated epitope on the spike and retains binding to the S-GSAS-Omicron and Delta spikes despite considerable differences in their S1 subunit dynamics. Two RBD-directed Abs, DH1044 and DH1193, that do not compete for ACE2 binding (*20*) retained binding to the Omicron S protein. The DH1044 epitope was mapped by NSEM to a region adjacent to the epitope of Ab S309, although shifted towards residue L452 making it susceptible to the L452R substitution in the Delta spike (Fig. 6, A and C) (*20*). Mapping of the DH1193 epitope by NSEM revealed an epitope in between the S309 and DH1044 epitopes (fig. S13).

We next probed the Omicron S2 subunit conformation by measuring binding to S2 targeting Abs (Fig. 6D and 7A, figs. S14 to S16). We have previously described the binding of HIV-1 neutralizing, Fab-dimerized glycan-reactive (FDG) Ab 2G12 to a quaternary glycan cluster in the S2 subunit of the SARS-CoV-2 S protein (*27*), and have demonstrated that 2G12 binding is sensitive to changes in S protein conformation (figs. S14 to S16) (*5, 23*). 2G12 and a panel of FDG Abs targeting the same glycan cluster (*27*) showed glycan-dependent binding to the Delta and Omicron S proteins (Fig. 6D and fig. S16). Binding of 2G12 to the Omicron S protein was weaker than its binding to the G614 and Delta S proteins, suggesting altered presentation of the glycan cluster, either due to a global change in S2 conformation or a local change due to the stabilization of glycan 709 by the D796Y substitution resulting in a change in presentation of the glycan epitope (Fig. 6D). These results are consistent with considerable Omicron S protein structural shifts and suggest that S2 dynamics and flexibility are impacted by its acquired mutations.

### Altered fusion peptide dynamics in the Omicron S protein

We next tested binding of the G614, Delta and Omicron spikes to SARS-CoV-2 fusion peptide (FP)-directed Ab DH1058 by ELISA (Fig. 7A) (*20*). DH1058 binds a 25-residue peptide spanning residues 808-833 that includes the FP (*5*), and showed ~6-fold increased binding to S-GSAS-Omicron compared to the G614 and Delta S proteins, suggesting greater access of DH1058 to the FP in the Omicron S protein (Fig. 7A). Binding rate and equilibrium constants (kon, koff and KD) measured by SPR revealed no differences between the variants tested (Fig. 7B). As the ELISA assay measures binding on a timescale slower than captured by the SPR assay, these results suggested time-dependent changes in the accessibility or presentation of the DH1058 epitope as the Ab is incubated with the spike for longer times in an ELISA experiment. The FP residues that are targeted by DH1058 were well-resolved in the cryo-EM reconstruction of the Omicron S protein (Fig. 7C) with more residues resolved in the cryo-EM reconstruction compared to other variant S proteins. The overall orientation of the FP was conserved between D614G, Alpha, Beta, Delta and Omicron S protein structures (Fig. 7C). While several attempts to obtain cryo-EM structures of DH1058 bound to furin-cleaved and uncleaved spikes were unsuccessful, we obtained a crystal structure of DH1058 Fab bound to a peptide comprising FP residues 808 to 833 at a resolution of 2.15 Å (P 1 21 1 space group) (Fig. 7D and table S2). The interaction between DH1058 and the FP is mediated by all heavy chain (HC) complementary determining regions (CDRs). The portion of the fusion peptide between residues 816 and 825 defines the interaction with Ser816 and Asp820 sidechains forming hydrogen bonds (H-bond) with the HCDR2 residues Tyr53, Glu54, Arg56 and Asn57 side chains. The HCDR1 Asp31 main chain carbonyl formed an H-bond with the FP Arg815 sidechain, while the HCDR3 Tyr115 and Tyr116 formed sidechain-to-sidechain H-bonds with Glu819 and Lys825 respectively (Fig. 7D). Aligning the structure of the trimeric pre-fusion Omicron S protein using the FP fragment from the crystal structure for superposition revealed clashes between the SD2 subdomain and bound DH1058 HC loops 13-17, 61-68 and 84-88, and the S2 subunit HR1 subdomain and DH1058 HCDR2 (Fig. 7E). These data show that the binding of DH1058 to the SARS-CoV-2 FP, as revealed by the crystal structure, is incompatible with the structures of the pre-fusion SARS-CoV-2 S proteins. Taken together, these data suggest that a weak initial contact of the DH1058 with the FP is made in the pre-fusion S protein ectodomains as captured in the SPR assay, followed by a conformational change in the spike leading to greater FP exposure and stable binding of the DH1058 Fab. DH1058 Fab cannot bind stably to the pre-fusion conformation of the SARS-CoV-2 S protein, consistent with its lack of SARS-CoV-2 neutralization activity (*20*) and would require greater exposure of the FP to make a stable interaction avoiding clashes with adjacent regions of the pre-fusion S protein. Our data here suggest that the conformational changes leading to greater FP exposure occur more readily in the Omicron S protein. Taken together, these results show altered flexibility and ease of exposure and release around the fusion peptide region in the S2 subunit of the Omicron spike relative to other variants.

**Figure 7.**
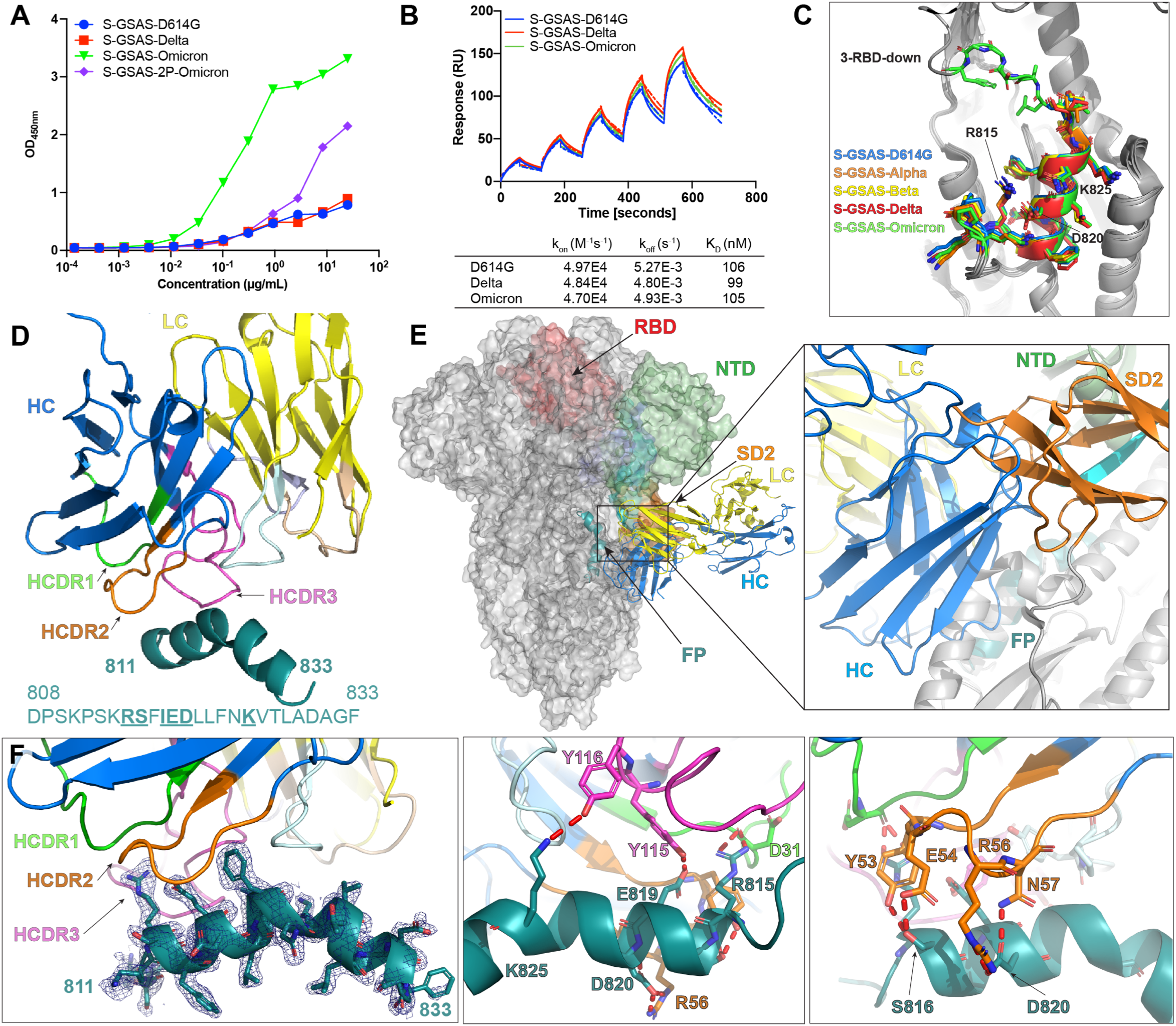
Binding of DH1058 to SARS-CoV-2 fusion peptide. (A) Binding of DH1058 to spike variants measured by ELISA. (B) Kinetics of DH1058 Fabs binding to spike variants measured by SPR. The full lines are the binding sensorgrams and the dotted lines show fits of the data to a 1:1 Langmuir binding model. The on-rate (k_on_, M-1s-1), off-rate (k_off_, s-1) and affinity (K_D_, nM) for each interaction are indicated in the table. (C) Conservation of the fusion peptide conformation in variant S protein structures. S-GSAS-D614G (PDB 7KDK, blue), S-GSAS-Alpha (B.1.1.7, PDB 7LWS, orange), S-GSAS-Beta (B.1.351, PDB 7LYM, yellow), S-GSAS-Delta (B.1.617.2, Red) and S-GSAS-Omicron (B.1.529, green) were superimposed in pymol using the extra fit method and residues 909-1036. (D) Crystal structure of DH1058 Fab variable region (heavy chain in blue and light chain in yellow) bound to a peptide comprising the SARS-CoV-2 S protein residues 808 to 833 (in teal). CDRs are colored green (HCDR1), orange (HCDR2), pink (HCDR3), wheat (LCDR1), pale purple(LCDR2) and pale blue (LCDR3).The bottom panel show the electron density of the peptide in the crystal structure. (E) Binding footprints of DH1058(colored in blue and yellow as in panel D) on SARS-CoV-2 S protein(PDB 7KDK).The S protein is shown as surface with 2 protomers colored gray and the protomer used for alignment with the crystal structure of DH1058-fusion peptide complex has its S1 subunits colored pale green (NTD -N-terminal domain), cyan (N2R -NTD-to-RBD linker), red (RBD-receptor-binding domain) and dark blue and orange (SD1 and SD2 respectively -subdomain 1 and 2). The inset panel shows a clash between DH1058 HC and the SARS-CoV-2 SD2 subdomain. A clash is also observed for between the HC and the S2 subunit HR1 subdomain (in grey). (F) Contacts (indicated by red dashed lines) between the DH1058 Fabs and the S2 peptide in the crystal structure.

## Discussion

As SARS-CoV-2 continues to evolve, the emergence of the Omicron variant is poised to change the course of the COVID-19 pandemic with its unprecedented transmissibility and immune evasion. The Omicron spike protein, that is central to defining these properties, is riddled with mutations in both its receptor binding S1 subunit, and its fusion subunit S2. As the exposure of Ab and receptor binding sites can be affected by both direct substitutions at the binding interface, and conformational masking of key sites, we have sought here to understand the conformational changes of the Omicron spike resulting from its altered primary sequence. Our structural studies were performed in our previously established platform S-GSAS that did not contain any extraneous stabilizing mutations in the S2 subunit, thus aiming to visualize S protein conformations in a more native format (*5, 12*). Indeed, we were able to resolve a more varied repertoire of structural states of the Omicron S protein than were revealed by several studies that have used constructs with stabilizing Proline mutations in the S2 subunit (*15–18*). The Omicron S protein presented a substantially different domain organization compared to other variants (Figure 1), differences that we were able to visualize in our structures and quantify using sets of intra- and inter-protomer vectors (Figures 1, 4 and 5). We found a tightly packed RBD-RBD interface in the 3-RBD-down state, with new interprotomer interactions mediated by a RBD loop harboring the S371L, S373P and S375F substitutions in one protomer, and a Y505H substitution in the adjacent interacting protomer. The close packing of the down state RBDs in Omicron is distinct from how the previous VOCs have evolved. While previous VOCs maximized transmissibility by favoring open-states of the spike and immune evasion by mutating common Ab epitopes, acquisition of RBD-down state stabilizing mutations is a significant change in a different direction. Stabilizing the RBD-down state bolsters immune evasion as it occludes highly immunogenic sites that bind very potent Abs. Moreover, the Omicron variant is more transmissible than any variant isolated so far. How does the Omicron spike achieve high transmissibility, which would require open states of the spike to engage receptor and undergo fusion, while also bolstering RBD-down state stability? The answer to this potentially lies in two aspects of our structural analysis. First, we showed that locking down the typically mobile RBDs results in a metastable spike with rearrangements in the critical peptide N2R that connects the NTD and RBD in a protomer, such that one protomer in 3-RBD-down spike is primed to transition to the up-state. Thus, the stabilization of the 3-RBD-down state is balanced by an enhanced propensity to adopt the up-state due to rearrangements in the N2R linker. Second, by combining binding assays, X-ray crystallography and cryo-EM, we uncovered evidence for altered plasticity of the fusion peptide in the Omicron spike compared to other variants, including Delta. Despite extensive stabilization of the Omicron spike, the functionally critical FP is more easily exposed in Omicron as measured by binding to a fusion peptide directed Ab. Thus, the increased transmissibility of the Omicron spike may be facilitated by a combined effect of the ease of accessing the RBD-up state despite stabilization of the down-state RBD-RBD interface, retained affinity for ACE2 interactions despite the large number of RBD mutations, as well as by more ready release of the FP. Close monitoring of the continued evolution of the structure of future variants on the Omicron template will be required to achieve a deep understanding of Omicron pathobiology, and to anticipate the immune escape potential of the further evolved variants.

## Materials and Methods

### Data Availability

Cryo-EM reconstructions and atomic models generated during this study have been deposited at wwPDB and EMBD (https://www.rcsb.org; http://emsearch.rutgers.edu) under the accession codes PDB IDs 7TF8, 7TL1, 7TEI, 7TL9, 7TGE, 7TOU, 7TOX, 7TOY, 7TOZ, 7TP0, 7TP1, 7TP2, 7TOV, 7TP7, 7TP8, 7TP9, 7TPA, 7TPC, 7TPE, 7TPF, 7TPH, 7TPL, 7TLA, 7TLB, 7TLC, 7TLD, 7THT and 7THE, and EMDB IDs 25865, 25983, 25846, 25984, 25880, 26038, 26040, 26041, 26042, 26043, 26045, 26046, 26039, 26047, 26048, 26049, 26050, 26051, 26052, 26053, 26055, 26059, 25985, 25986, 25987, 25988, 25904 and 25893. The crystal structure of DH1058 Fab bound to the SARS-CoV-2 fusion peptide is deposited at wwPDB with PDB ID 7TOW. Vector analysis scripts are available at: https://doi.org/10.5281/zenodo.4926233.

### Plasmids

Mutagenesis of all plasmids generated by this study were performed and the sequence confirmed by GeneImmune Biotechnology (Rockville, MD). The SARS-CoV-2 spike protein ectodomain constructs comprised the S protein residues 1 to 1208 (GenBank: MN908947) with the D614G mutation, the furin cleavage site (RRAR; residue 682-685) mutated to GSAS, a C-terminal T4 fibritin trimerization motif, a C-terminal HRV3C protease cleavage site, a TwinStrepTag and an 8XHisTag. All spike ectodomains were cloned into the mammalian expression vector pαH and have been deposited to Addgene (*10*) (https://www.addgene.org).

### Cell culture and protein expression

GIBCO FreeStyle 293-F cells (embryonal, human kidney) were maintained at 37°C and 9% CO_2_ in a 75% humidified atmosphere in FreeStyle 293 Expression Medium (GIBCO). Plasmids were transiently transfected using Turbo293 (SpeedBiosystems) and incubated at 37°C, 9% CO2, 75% humidity with agitation at 120 rpm for 6 days. On the day following transfection, HyClone CDM4HEK293 media (Cytiva, MA) was added to the cells. Antibodies were produced in Expi293F cells (embryonal, human kidney, GIBCO). Cells were maintained in Expi293 Expression Medium (GIBCO) at 37°C, 120 rpm and 8% CO2 and 75% humidity. Plasmids were transiently transfected using the ExpiFectamine 293 Transfection Kit and protocol (GIBCO).

### Protein purification

On the 6^th^ day post transfection, spike ectodomains were harvested from the concentrated supernatant. The spike ectodomains were purified using StrepTactin resin (IBA LifeSciences) and size exclusion chromatography (SEC) using a Superose 6 10/300 GL Increase column (Cytiva, MA) equilibrated in 2mM Tris, pH 8.0, 200 mM NaCl, 0.02% NaN3. All steps of the purification were performed at room temperature and in a single day. Protein quality was assessed by SDS-PAGE using NuPage 4-12% (Invitrogen, CA). The purified proteins were flash frozen and stored at −80°C in single-use aliquots. Each aliquot was thawed by a 20-minute incubation at 37 °C before use. Antibodies were purified by Protein A affinity and digested to their Fab state using LysC. ACE2 with human Fc tag was purified by Protein A affinity chromatography and SEC.

### SPR

Antibody binding to SARS-CoV-2 spike was assessed using SPR on a Biacore T-200 (Cytiva, MA, formerly GE Healthcare) with HBS buffer supplemented with 3 mM EDTA and 0.05% surfactant P-20 (HBS-EP+, Cytiva, MA). All binding assays were performed at 25 °C. Spike variants were captured on a Series S Strepavidin (SA) chip (Cytiva, MA) coated at 100 nM (60s at 10μL/min). Fabs were injected at concentrations ranging from 0.625 nM to 800 nM (prepared in a 2-fold serial dilution manner) over the S proteins using the single cycle kinetics mode with 5 concentrations per cycle. The surface was regenerated after the last injection with 3 pulses of a 50mM NaoH + 1M NaCl solution for 10 seconds at 100μL/min. Sensogram data were analyzed using the BiaEvaluation software (Cytiva, MA)

### Negative-stain electron microscopy

Fab-spike complex of DH1044 was generated by mixing 5.9 μg of spike with 6.2 μg of Fab in ~25 μl of HEPES-buffered saline (HBS) containing 20 mM HEPES and 150 mM NaCl, pH 7.4, and incubating at 37 °C for 1 hr and using sample directly for negative stain without further purification. Fab-spike complex of DH1193 was generated by mixing 20 μg of spike with 28 μg of Fab in ~200 μl of phosphate-buffered saline and incubating 1 hr at 37 °C. Sample was then brought to room temperature and diluted with 800 μl HBS, mixed, and then diluted with HBS augmented with 16 mM glutaraldehyde (Electron Microscopy Sciences, PA), fixed for 5 min, quenched by addition of 40 μl of 1 M Tris buffer, pH 7.4, and concentrated in a 2-ml 100-kDa MCWO Amicon centrifugal concentrator by spinning 10 min at 4000 rpm in a Sorvall benchtop centrifuge with a swinging-bucket rotor, yielding a final volume of ~75 μl. Protein concentration was measured using a Nanodrop which reported a nominal spike concentration of 0.6 mg/ml

For negative stain, samples were diluted to 0.1 mg/ml with HBS augmented with 5 g/dl glycerol and 8 mM glutaraldehyde. After 5 min incubation, excess glutaraldehyde was quenched by adding sufficient 1 M Tris stock, pH 7.4, to give 75 mM final Tris concentration and incubated for 5 min. Quenched sample was applied to a glow-discharged carbon-coated EM grid (Electron Microscopy Sciences, PA, CF300-Cu) for 10-12 second, then blotted, and stained with 2 g/dL uranyl formate (Electron Microscopy Sciences, PA), for 1 min, blotted and air-dried. Grids were examined on a Philips EM420 electron microscope operating at 120 kV and images were collected at a nominal magnification of 82,000x on a 4 Mpix CCD camera at 4.02 Å/pixel for DH1044 data, or at 49,000x on a 76 Mpix CCD camera at 2.4 Å/pixel for DH1193 data. Images were analyzed and 3D reconstructions generated using standard protocols with Relion 3.0 (*28*).

### ELISA assays

Spike ectodomains tested for antibody- or ACE2-binding in ELISA assays as previously described. Assays were run in two formats *i.e*., antibodies/ACE2 coated, or spike coated. For the first format, the assay was performed on 384-well plates coated at 2 μg/ml overnight at 4°C, washed, blocked and followed by two-fold serially diluted spike protein starting at 25 μg/mL. Binding was detected with polyclonal anti-SARS-CoV-2 spike rabbit serum (developed in our lab), followed by goat anti-rabbit-HRP (Abcam, Ab97080) and TMB substrate (Sera Care Life Sciences, MA). Absorbance was read at 450 nm. In the second format, serially diluted spike protein was bound in wells of a 384-well plates, which were previously coated with streptavidin (Thermo Fisher Scientific, MA) at 2 μg/mL and blocked. Proteins were incubated at room temperature for 1 hour, washed, then human mAbs were added at 10 μg/ml. Antibodies were incubated at room temperature for 1 hour, washed and binding detected with goat anti-human-HRP (Jackson ImmunoResearch Laboratories, PA) and TMB substrate.

Recombinant FDG mAbs were tested for binding to the SARS-CoV-2 Omicron spike (S-GSAS-Omicron, Lot: 486KJ), Delta spike (S-GSAS/B.1.617.2.v1, Lot: 076XH), SARS-CoV-2 spike (nCoV-1_2 ProRev+D614G, Lot: 001AM), in ELISA in the absence or presence of single monomer D-mannose as previously described (PMID: 34019795). Briefly, spike proteins (20ng) were captured by streptavidin (30ng per well) to individual wells of a 384-well Nunc-absorb ELISA plates using PBS-based buffers and assay conditions as previously described (PMID: 34019795; PMID: 28298421; PMID: 28298420). Commercially obtained D-mannose (Sigma, St. Louis, MO) was used to outcompete mAb binding to glycans on the spike proteins; D-mannose solutions were also produced in ELISA PBS-based glycan buffers at a concentration of [1M] D-mannose as described (PMID: 34019795). Mouse anti-monkey IgG-HRP (Southern Biotech, CAT# 4700-05) and Goat anti-human IgG-HRP (Jackson ImmunoResearch Laboratories, CAT# 109-035-098) secondary antibodies were used to detect antibody bound to the spike proteins. HRP detection was subsequently quantified with 3,30,5,50-tetramethylbenzidine (TMB) by measuring binding levels at an absorbance of 450nm, and binding titers were also reported as Log area under the curve (AUC).

### Cryo-EM

Purified SARS-CoV-2 spike ectodomains were diluted to a concentration of ~1.5 mg/mL in 2 mM Tris pH 8.0, 200 mM NaCl and 0.02% NaN3 and 0.5% glycerol was added. A 2.3-μL drop of protein was deposited on a Quantifoil-1.2/1.3 grid (Electron Microscopy Sciences, PA) that had been glow discharged for 10 seconds using a PELCO easiGlow™ Glow Discharge Cleaning System. After a 30-second incubation in >95% humidity, excess protein was blotted away for 2.5 seconds before being plunge frozen into liquid ethane using a Leica EM GP2 plunge freezer (Leica Microsystems). Frozen grids were imaged using a Titan Krios (Thermo Fisher) equipped with a K3 detector (Gatan). The cryoSPARC (*29*) software was used for data processing. Phenix (*30, 31*), Coot (*32*), Pymol (*33*), Chimera (*34*), ChimeraX (*35*) and Isolde (*36*) were used for model building and refinement.

### Vector Based Structure Analysis

Vector analysis of intraprotomer domain positions was performed as described previously using the Visual Molecular Dynamics (VMD) (*37*) software package Tcl interface. For each protomer of each structure, Cα centroids were determined for the NTD (residues 27 to 69, 80 to 130, 168 to 172, 187 to 209, 216 to 242, and 263 to 271), NTD′ (residues 44 to 53 and 272 to 293), RBD (residues 334 to 378, 389 to 443, and 503 to 521), SD1 (residues 323 to 329 and 529 to 590), SD2 (residues 294 to 322, 591 to 620, 641 to 691, and 692 to 696), CD (residues 711 to 716 1072 to 1121), and a S2 sheet motif (S2s; residues 717 to 727 and 1047 to 1071). Additional centroids for the NTD (NTD_c_; residues 116 to 129 and 169 to 172) and RBD (RBD_c_; residues 403 to 410) were determined for use as reference points for monitoring the relative NTD and RBD orientations to the NTD′ and SD1, respectively. Vectors were calculated between the following within protomer centroids: NTD to NTD′, NTD′ to SD2, SD2 to SD1, SD2 to CD, SD1 to RBD, CD to S2s, NTD_c_ to NTD, RBD to RBD_c_. Vector magnitudes, angles, and dihedrals were determined from these vectors and centroids. Inter-protomer domain vector calculations for the SD2, SD1, and NTD′ used these centroids in addition to anchor residue Cα positions for each domain including SD2 residue 671 (SD2a), SD1 residue 575 (SD1a), and NTD′ residue 276 (NTD′a). These were selected based upon visualization of position variation in all protomers used in this analysis via alignment of all of each domain in PyMol (*33*). Vectors were calculated for the following: NTD′ to NTD′_r_, NTD′ to SD2, SD2 to SD2_r_, SD2 to SD1, SD1 to SD1_r_, and SD1 to NTD′. Angles and dihedrals were determined from these vectors and centroids. Vectors for the RBD to adjacent RBD and RBD to adjacent NTD were calculated using the above RBD, NTD, and RBD_c_ centroids. Vectors were calculated for the following: RBD_2_ to RBD_1_, RBD_3_ to RBD_2_, and RBD_3_ to RBD_1_. Angles and dihedrals were determined from these vectors and centroids. Principal components analysis, K-means clustering, and Pearson correlation (confidence interval 0.95, p<0.05) analysis of vectors sets was performed in R(*38*). Data were centered and scaled for the PCA analyses. Principal components analysis, K-means clustering, and Pearson correlation (confidence interval 0.95, p < 0.05) analysis of vectors sets was performed in R. Data were centered and scaled for the PCA analyses.

### Difference distance matrices (DDM)

DDM were generated using the Bio3D package (*39*) implemented in R (R Core Team (2014). R: A language and environment for statistical computing. R Foundation for Statistical Computing, Vienna, Austria. URL http://www.R-project.org/)

### Crystallization and determination of the structure of DH1058 fab bound to a fusion peptide fragment

After size exclusion purification, DH1058 fab was concentrated to 26 mg/mL. The fusion peptide fragment was solubilized in PBS + 10% DMSO at a concentration of 10 mg/mL. The protein and the peptide were mixed at a Fab:peptide molar ratio of 1:2. Crystals were grown in 20% PEG3000, 100mM Tris base/HCl pH 7.0, 200mM calcium acetate at 22°C in a sitting drop vapor diffusion setting using a drop ratio of 0.4 μL protein : 0.2 μL reservoir solution. Large UV-active plate shaped crystals were observed after 24 hours. A single crystal was cryopreserved directly from the drop. Diffraction data was collected at the Advanced Photon Source using sector 22ID beamline. The collected diffraction images were indexed, integrated, and scaled using HKL2000 (*40*) Initial phases were calculated by molecular replacement using Phenix.PHASER (*41, 42*) and the PDB 5GGU (Crystal structure of tremelimumab Fab) as a search model. Iterative rounds of manual model building using Coot (*32*) and automatic refinement in PHENIX (*30, 31*) were performed. Data collection and refinement statistics are summarized in **Table S2**. The refined structure has been deposited to the Protein Data Bank (http://www.pdb.org) under the accession code 7TOW.

## Acknowledgements

Cryo-EM data were collected at the Duke Krios at the Duke University Shared Materials Instrumentation Facility (SMIF), a member of the North Carolina Research Triangle Nanotechnology Network (RTNN), which is supported by the National Science Foundation (award number ECCS-2025064) as part of the National Nanotechnology Coordinated Infrastructure (NNCI), and at the National Center for Cryo-EM Access and Training (NCCAT) and the Simons Electron Microscopy Center located at the New York Structural Biology Center, supported by the NIH Common Fund Transformative High Resolution Cryo-Electron Microscopy program (U24 GM129539) and by grants from the Simons Foundation (SF349247) and NY State. We thank Ed Eng, Daija Bobe and Nilakshee Bhattacharya for microscope alignments and assistance with cryo-EM data collection. This study utilized the computational resources offered by Duke Research Computing (http://rc.duke.edu; NIH 1S10OD018164-01) at Duke University. Use of the Advanced Photon Source was supported by the U. S. Department of Energy, Office of Science, Office of Basic Energy Sciences, under Contract No. W-31-109-Eng-38.

## Author contributions

S.M-C.G., R.H., V.S., A.M., K.Manne and P.A. determined and analyzed cryo-EM structures. S.M-C.G. determined and analyzed the X-ray crystallography structure and performed SPR experiments. S.M-C.G., R.C.H, V.S. and P.A. wrote the manuscript with help from all authors. V.S. K.J., X.H., A.M., M.S. and E.B. expressed and purified proteins. R.P., M.B., M.D., and M.M. performed ELISA assays. K.Mansouri and R.J.E. performed NSEM analysis. D.L., G.D.S., K.W., and W.W. analyzed data and edited the manuscript. K.O.S. provided key reagents. B.K. supervised variant sequences. B.F.H provided S RBD and S2 antibodies, supervised ELISA assays and edited the paper. P.A. supervised the study and reviewed all data.

