## Supplementary Data for "Structural diversity of the SARS-CoV-2 Omicron spike"

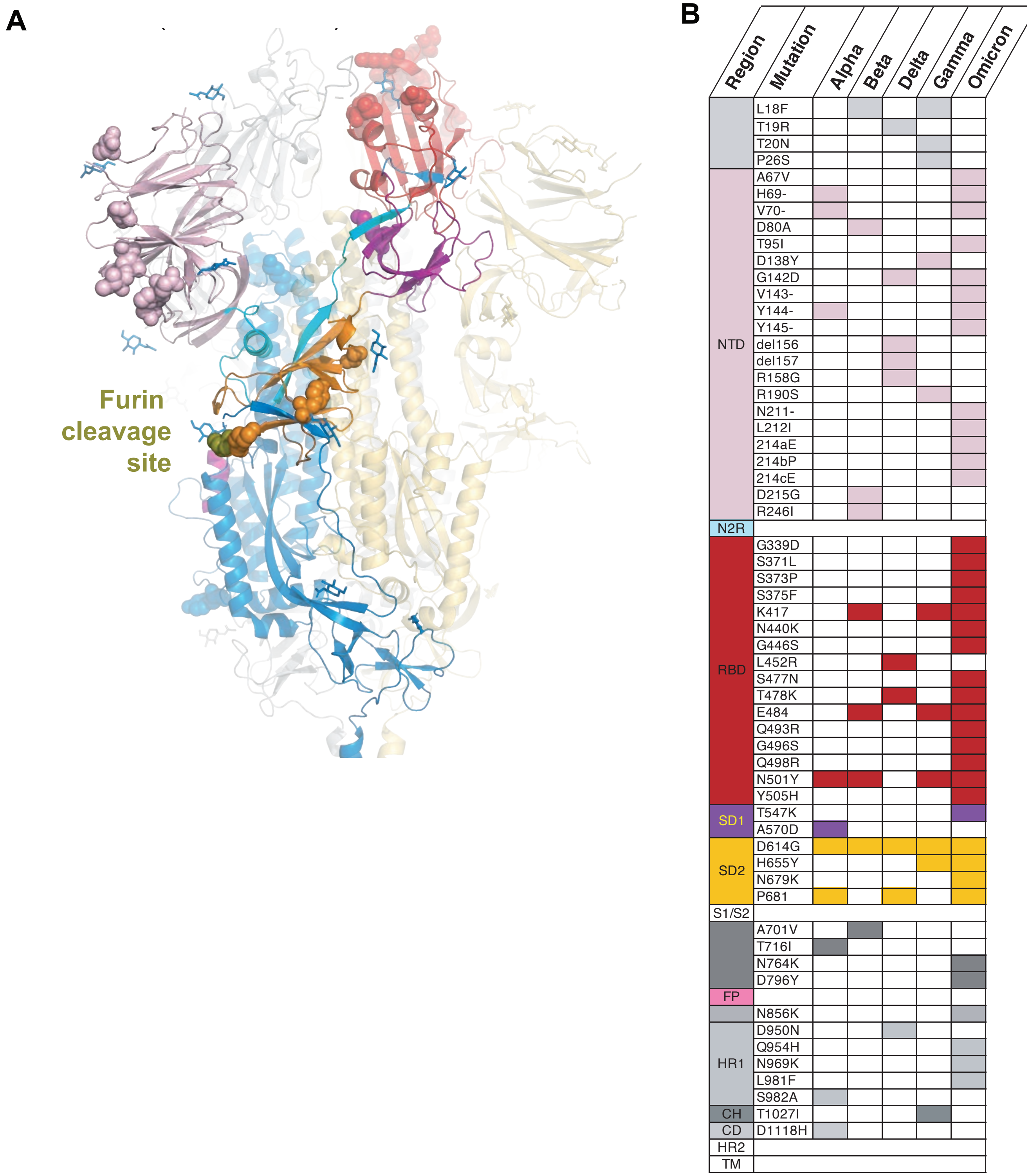
A
Furin cleavage site
B
Region
Mutation
Alpha
Beta
Delta
Gamma
Omicron
L18F
T19R
T20N
P26S
NTD
A67V
H69-
V70-
D80A
T95I
D138Y
G142D
V143-
Y144-
Y145-
del156
del157
R158G
R190S
N211-
L212I
214aE
214bP
214cE
D215G
R246I
N2R
RBD
G339D
S371L
S373P
S375F
K417
N440K
G446S
L452R
S477N
T478K
E484
Q493R
G496S
Q498R
N501Y
Y505H
SD1
T547K
A570D
SD2
D614G
H655Y
N679K
P681
S1/S2
A701V
T716I
N764K
D796Y
FP
N856K
HR1
D950N
Q954H
N969K
L981F
S982A
CH
T1027I
CD
D1118H
HR2
TM

**Fig. S1. SARS-CoV-2 spike (S) protein ectodomains of variants of concern (VOCs). (A)** S-GSAS-Omicron (B.1.1.529) variant model with mutations represented by spheres with comparison to the ancestral SARS-CoV-2 S protein in the 3-down state. The NTD (N-terminal domain, pale pink), N2R (NTD-to-RBD linker, cyan), RBD (receptor binding domain, red), and SD1 and SD2 (subdomains 1 and 2, purple and orange). The S2 subunit contains the FP (fusion peptide, magenta), and the furin cleavage site (deep olive). **(B)** Point mutation comparison of alpha, beta, delta, gamma, and omicron variants. The colored block corresponds with that variant holding the labeled mutation.Colors match the structure in A, with the following exceptions: HR1 (heptad repeat), CH (central helix), CD (connector domain), and HR2 (heptad repeat 2) regions.

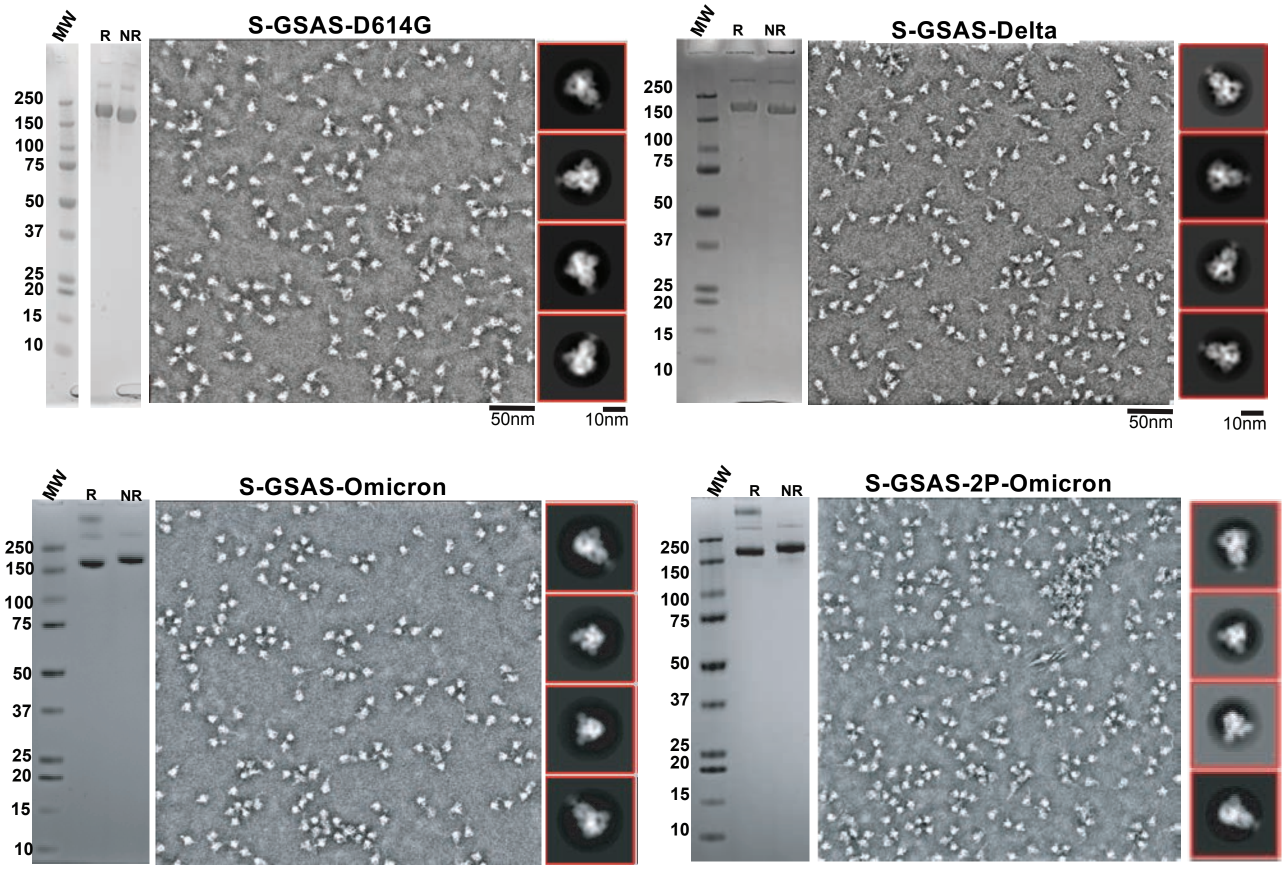


**Fig. S2. Quality control assessment of the SARS-CoV-2 spike variants preparations.**
SDS-PAGE of the purified SARS-CoV-2 S protein ectodomains, labels represent: R (Reduced) and NR (Non-Reduced), and representative NSEM micrographs and 2D class averages.

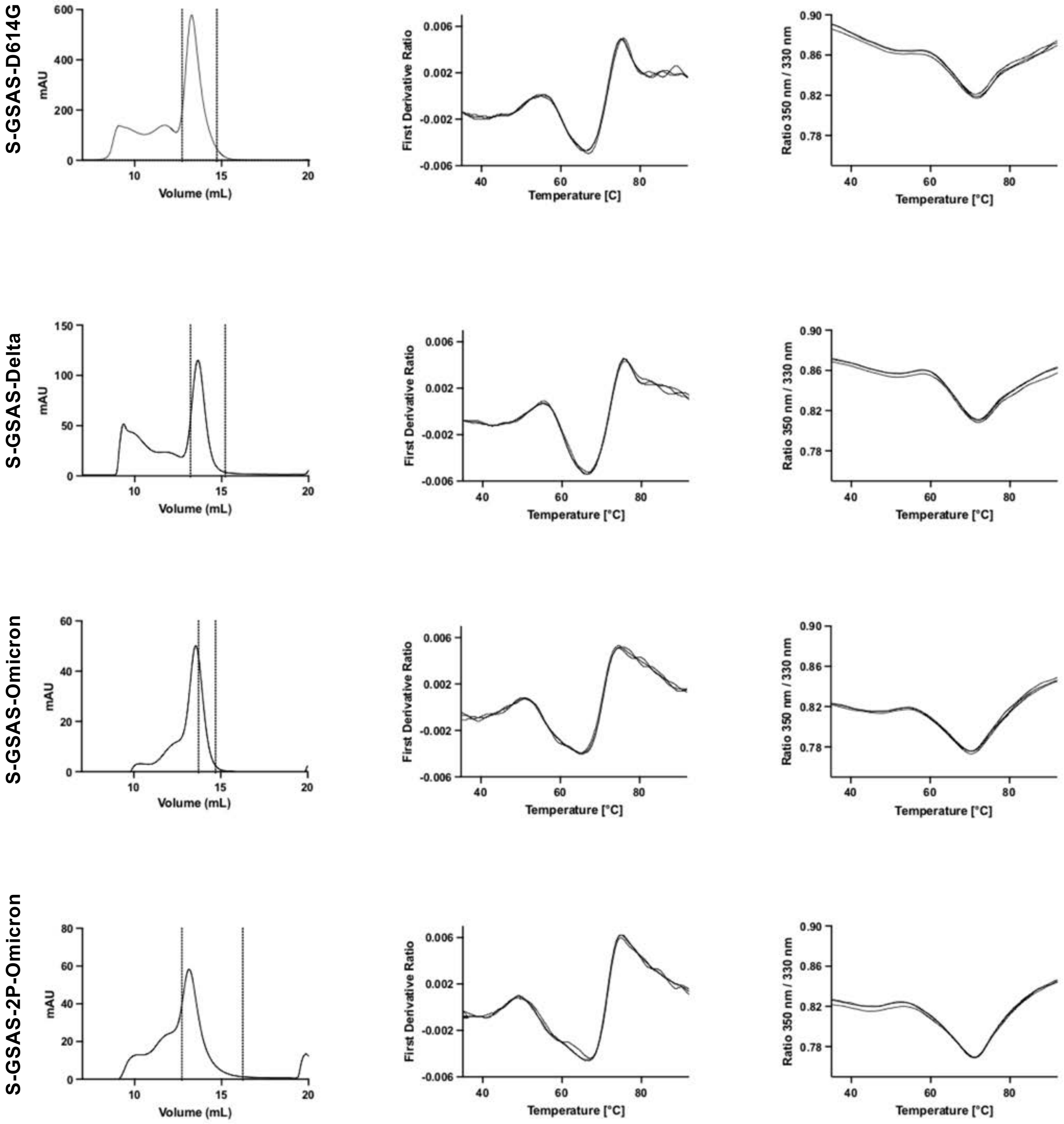


**Fig. S2 (contd). Quality control assessment of the spike variants preparations.**
From left to right, Size Exclusion Chromatography (SEC) profiles with vertical, dotted lines representing fractions that were pooled for subsequent studies; Differential Scanning Fluorimetry (DSF) first derivate ratio profile, and DSF smoothed profile of spike ectodomains.

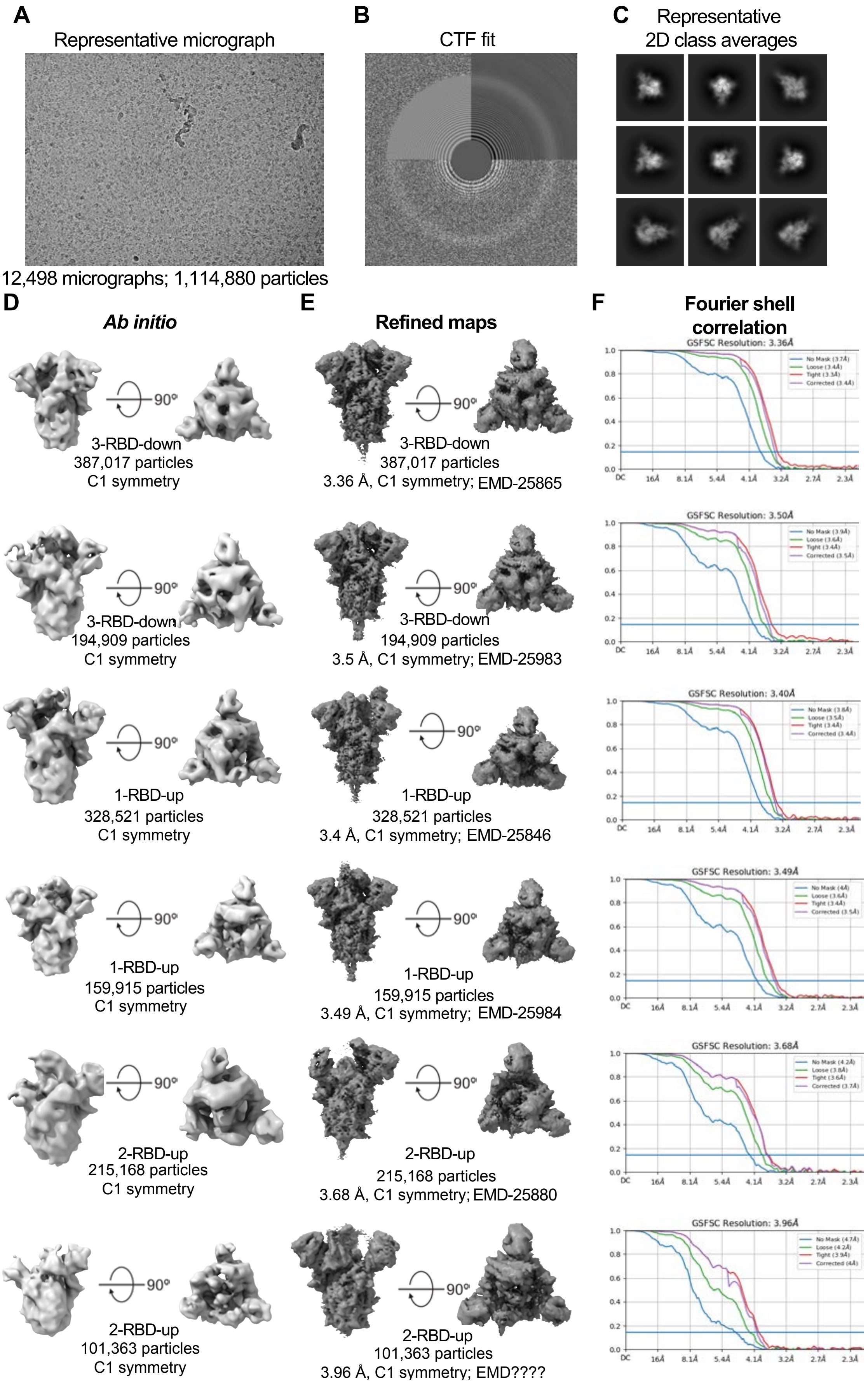
A
Representative micrograph
12,498 micrographs; 1,114,880 particles
B
CTF fit
C
Representative
2D class averages
D
Ab initio
E
Refined maps
F
Fourier shell
correlation
90°
3-RBD-down
387,017 particles
C1 symmetry
3-RBD-down
387,017 particles
3.36 Å, C1 symmetry; EMD-25865
GSFSC Resolution: 3.36Å
3-RBD-down
194,909 particles
C1 symmetry
3-RBD-down
194,909 particles
3.5 Å, C1 symmetry; EMD-25983
GSFSC Resolution: 3.50Å
1-RBD-up
328,521 particles
C1 symmetry
1-RBD-up
328,521 particles
3.4 Å, C1 symmetry; EMD-25846
GSFSC Resolution: 3.40Å
1-RBD-up
159,915 particles
C1 symmetry
1-RBD-up
159,915 particles
3.49 Å, C1 symmetry; EMD-25984
GSFSC Resolution: 3.49Å
2-RBD-up
215,168 particles
C1 symmetry
2-RBD-up
215,168 particles
3.68 Å, C1 symmetry; EMD-25880
GSFSC Resolution: 3.68Å
2-RBD-up
101,363 particles
C1 symmetry
2-RBD-up
101,363 particles
3.96 Å, C1 symmetry; EMD????
GSFSC Resolution: 3.96Å

**Fig. S3. Cryo-EM data processing for the S-GSAS-Omicron SARS-COV-2 S ectodomain.** **(A**) Representative micrograph. **(B)** CTF Fit. **(C)** Representative 2D class averages from cryo-EM dataset. **(D)** *Ab initio* reconstructions for the cryo-EM 3-RBD-down, 1-RBD-up, and 2-RBD-up states. **(E)** Refined maps for the cryo-EM corresponding states. **(F)** Fourier shell correlation curves for the cryo-EM corresponding states.

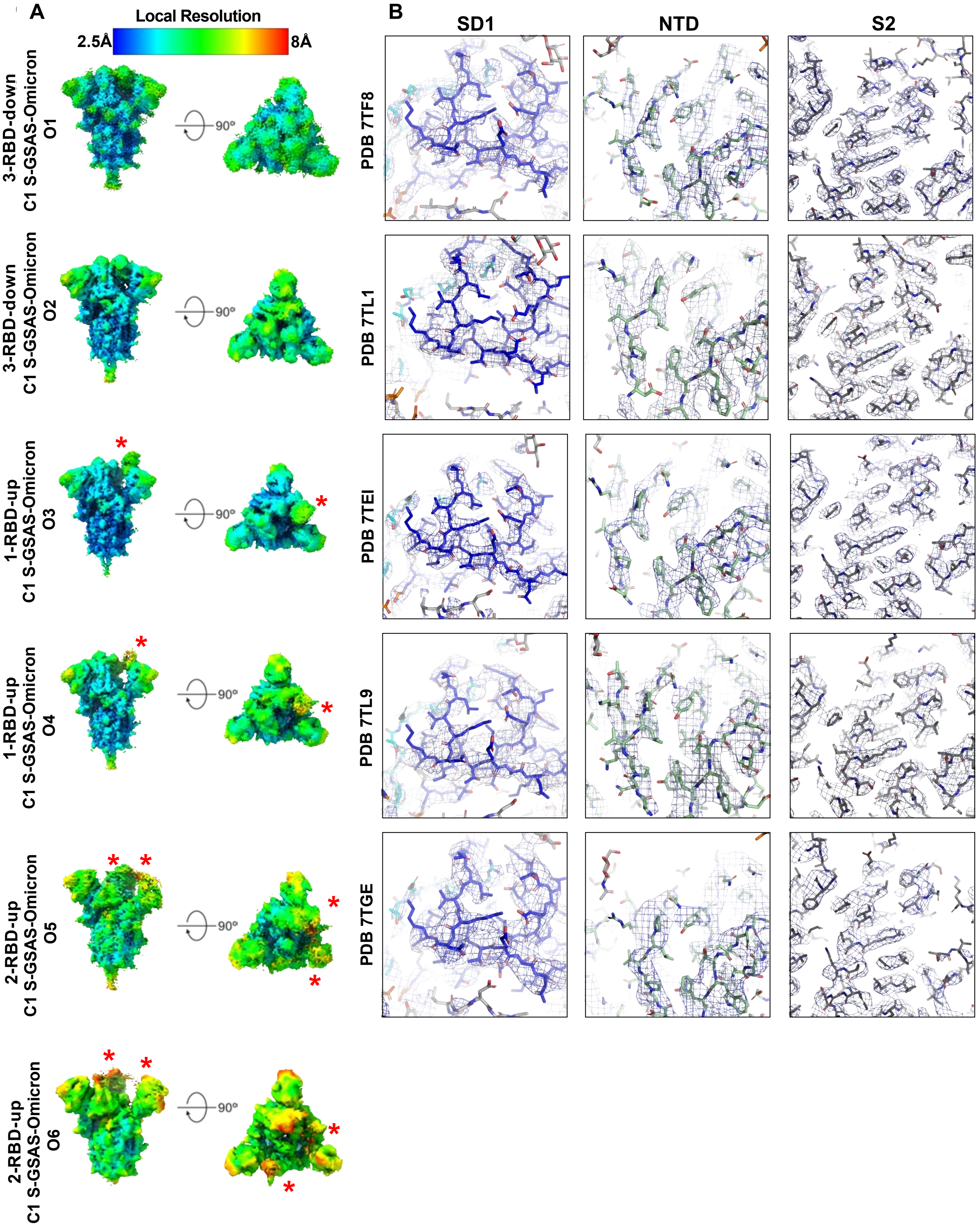


**Fig. S4. Quality assessment of the cryo-EM map and model fitting.**

**(A)** Refined maps colored by local resolution ranging from 2.5Å to 8Å. **(B)** Zoomed in views of SD1, NTD, and S2.

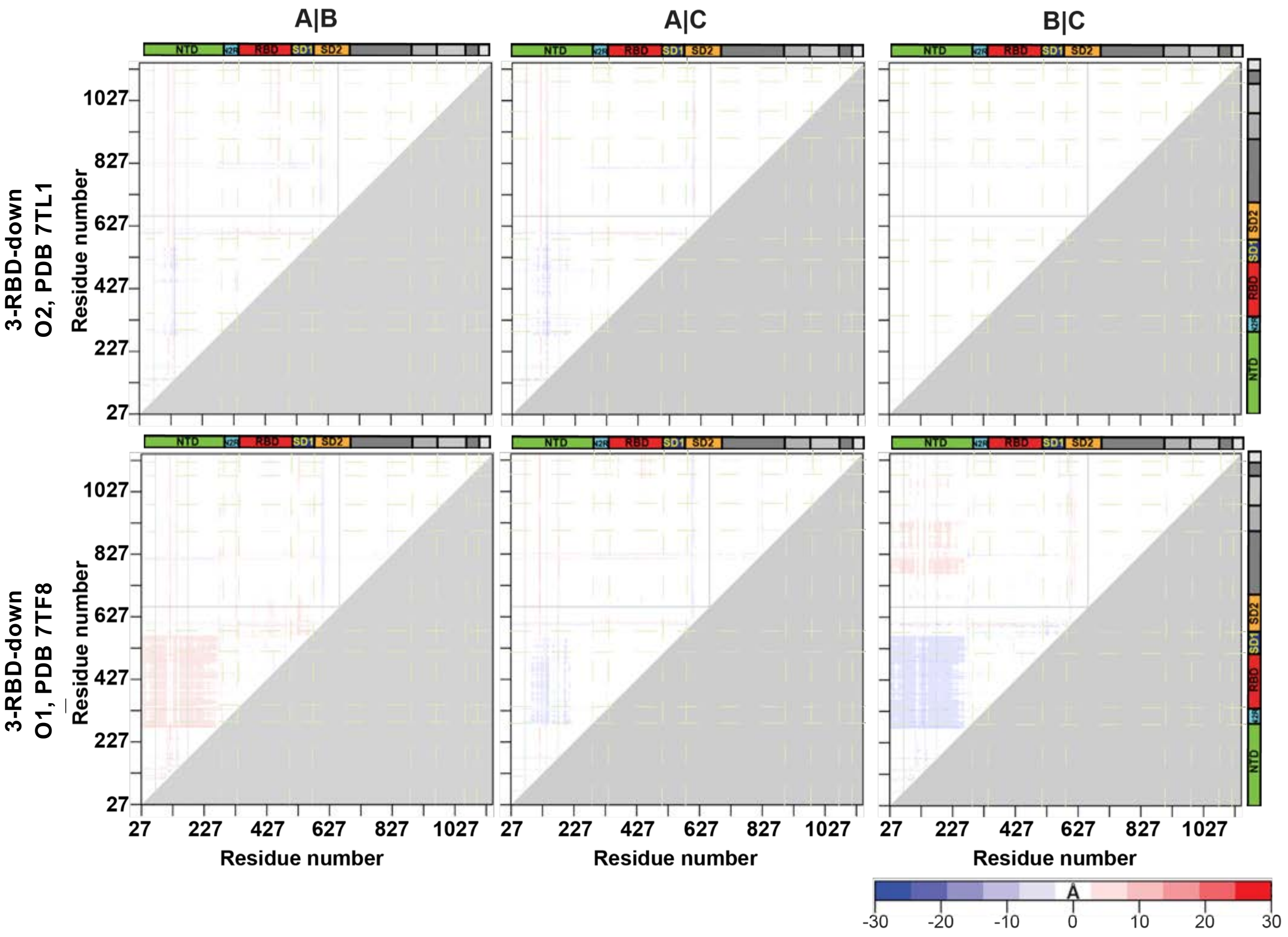


**Figure S5. Variability between protomers in the S-GSAS-Omicron S ectodomain 3-RBD-down structures.** DDM of the 3-RBD down structures comparing each chain of a structure to the other chains of the same structure. The coloring scale used for the DDM analysis is represented on the bottom right corner.

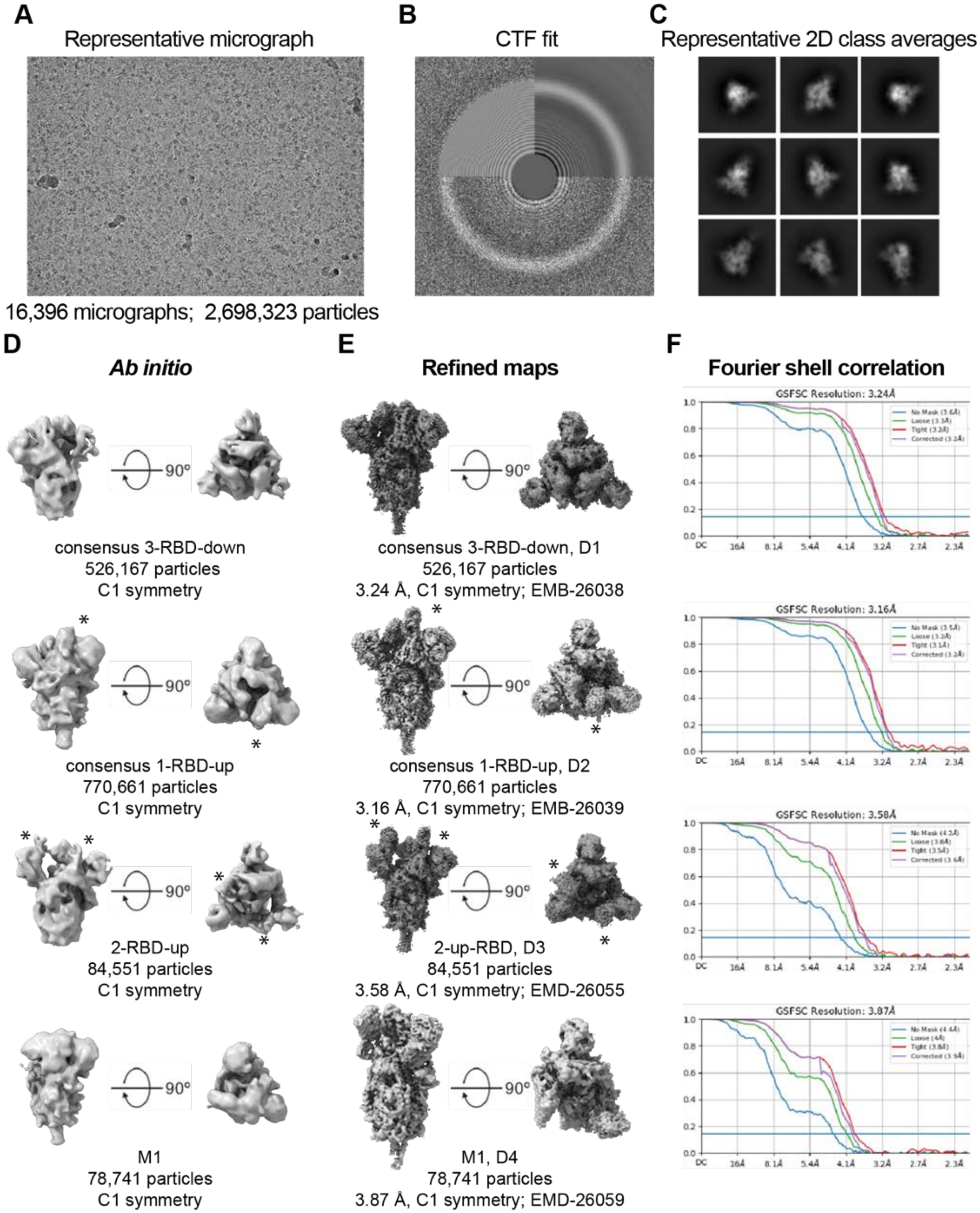
A
Representative micrograph
16,396 micrographs; 2,698,323 particles
B
CTF fit
C
Representative 2D class averages
D
Ab initio
90°
consensus 3-RBD-down
526,167 particles
C1 symmetry
consensus 1-RBD-up
770,661 particles
C1 symmetry
2-RBD-up
84,551 particles
C1 symmetry
M1
78,741 particles
C1 symmetry
E
Refined maps
consensus 3-RBD-down, D1
526,167 particles
3.24 Å, C1 symmetry; EMB-26038
consensus 1-RBD-up, D2
770,661 particles
3.16 Å, C1 symmetry; EMB-26039
2-up-RBD, D3
84,551 particles
3.58 Å, C1 symmetry; EMD-26055
M1, D4
78,741 particles
3.87 Å, C1 symmetry; EMD-26059
F
Fourier shell correlation
GSFSC Resolution: 3.24Å
GSFSC Resolution: 3.16Å
GSFSC Resolution: 3.58Å
GSFSC Resolution: 3.87Å
DC
16Å
8.1Å
5.4Å
4.1Å
3.2Å
2.7Å
2.3Å

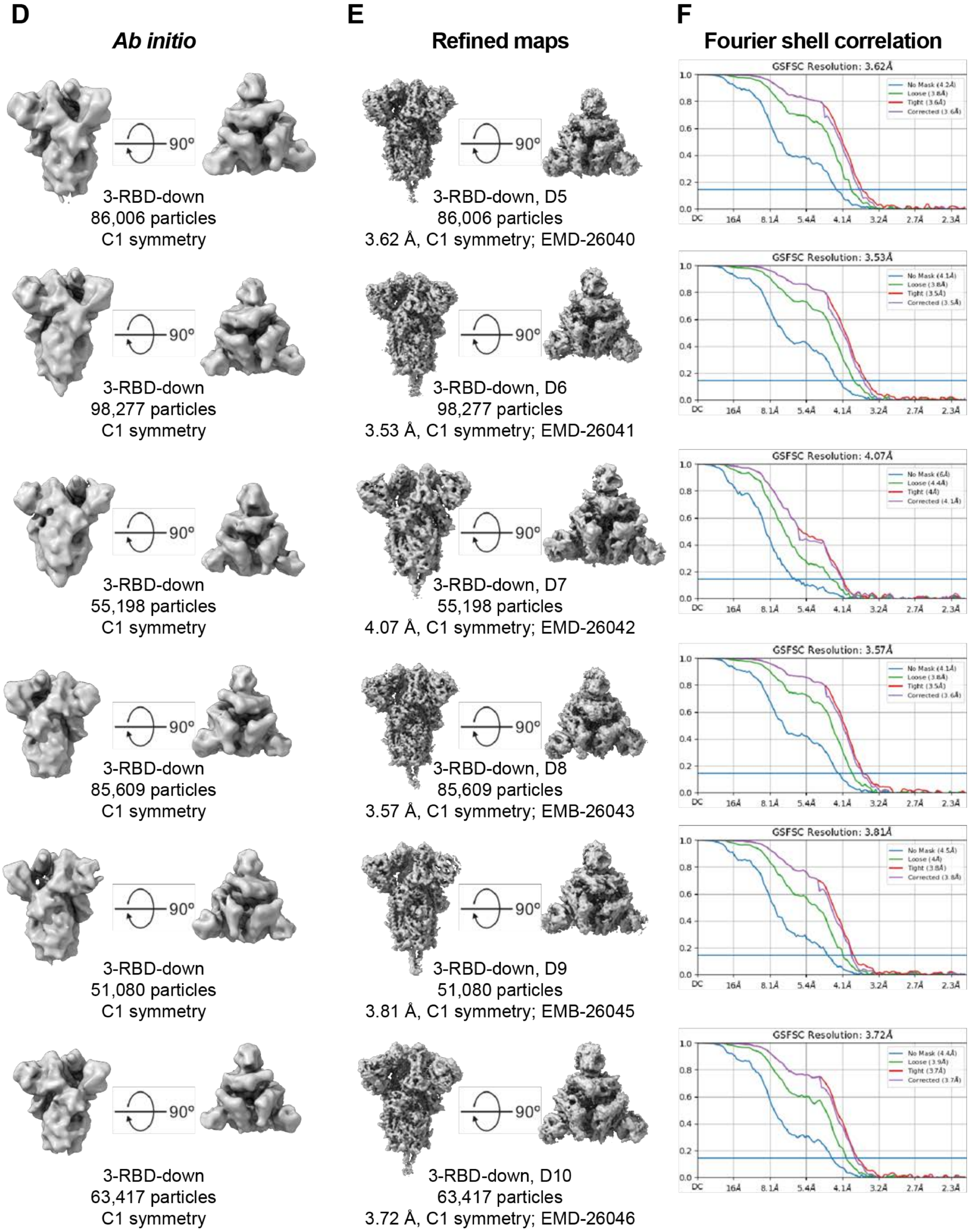
D
Ab initio
E
Refined maps
F
Fourier shell correlation
90°
3-RBD-down
86,006 particles
C1 symmetry
3-RBD-down, D5
86,006 particles
3.62 Å, C1 symmetry; EMD-26040
GSFSC Resolution: 3.62Å
3-RBD-down
98,277 particles
C1 symmetry
3-RBD-down, D6
98,277 particles
3.53 Å, C1 symmetry; EMD-26041
GSFSC Resolution: 3.53Å
3-RBD-down
55,198 particles
C1 symmetry
3-RBD-down, D7
55,198 particles
4.07 Å, C1 symmetry; EMD-26042
GSFSC Resolution: 4.07Å
3-RBD-down
85,609 particles
C1 symmetry
3-RBD-down, D8
85,609 particles
3.57 Å, C1 symmetry; EMB-26043
GSFSC Resolution: 3.57Å
3-RBD-down
51,080 particles
C1 symmetry
3-RBD-down, D9
51,080 particles
3.81 Å, C1 symmetry; EMB-26045
GSFSC Resolution: 3.81Å
3-RBD-down
63,417 particles
C1 symmetry
3-RBD-down, D10
63,417 particles
3.72 Å, C1 symmetry; EMD-26046
GSFSC Resolution: 3.72Å

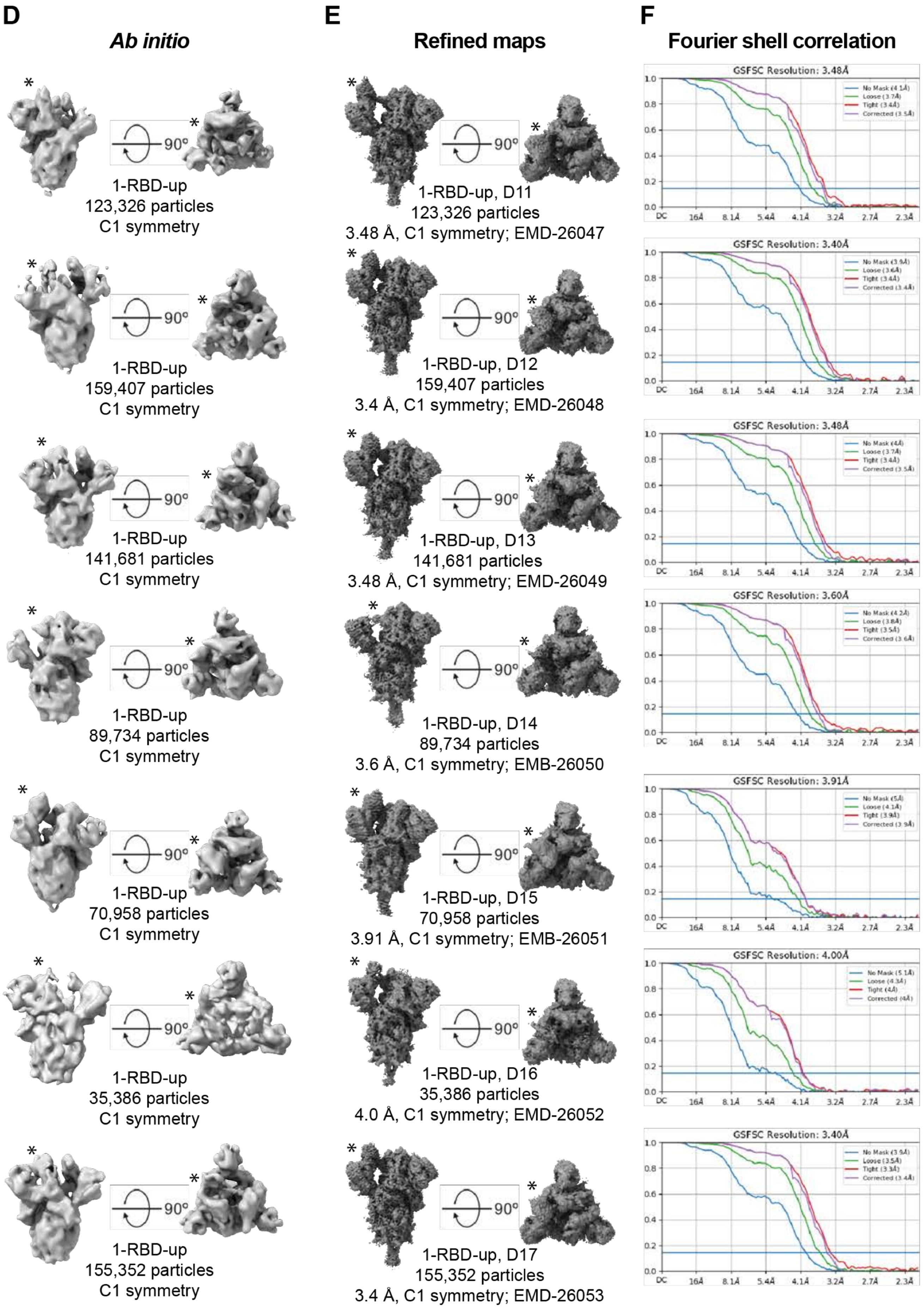
D
Ab initio
E
Refined maps
F
Fourier shell correlation
1-RBD-up
123,326 particles
C1 symmetry
1-RBD-up, D11
123,326 particles
3.48 Å, C1 symmetry; EMD-26047
GSFSC Resolution: 3.48Å
1-RBD-up
159,407 particles
C1 symmetry
1-RBD-up, D12
159,407 particles
3.4 Å, C1 symmetry; EMD-26048
GSFSC Resolution: 3.40Å
1-RBD-up
141,681 particles
C1 symmetry
1-RBD-up, D13
141,681 particles
3.48 Å, C1 symmetry; EMD-26049
GSFSC Resolution: 3.48Å
1-RBD-up
89,734 particles
C1 symmetry
1-RBD-up, D14
89,734 particles
3.6 Å, C1 symmetry; EMB-26050
GSFSC Resolution: 3.60Å
1-RBD-up
70,958 particles
C1 symmetry
1-RBD-up, D15
70,958 particles
3.91 Å, C1 symmetry; EMB-26051
GSFSC Resolution: 3.91Å
1-RBD-up
35,386 particles
C1 symmetry
1-RBD-up, D16
35,386 particles
4.0 Å, C1 symmetry; EMD-26052
GSFSC Resolution: 4.00Å
1-RBD-up
155,352 particles
C1 symmetry
1-RBD-up, D17
155,352 particles
3.4 Å, C1 symmetry; EMD-26053
GSFSC Resolution: 3.40Å
90º

**Fig. S6. Cryo-EM data processing for the S-GSAS-Delta SARS-COV-2 S ectodomain.** (**A**) Representative micrograph. **(B)** CTF Fit. **(C)** Representative 2D class averages from Cryo-EM dataset. **(D)** *Ab initio* reconstructions for the cryo-EM 3-RBD-down, 1-RBD-up, and 2-RBD-up states. **(E)** Refined maps for the cryo-EM corresponding states. **(F)** Fourier shell correlation curves for the cryo-EM corresponding states.

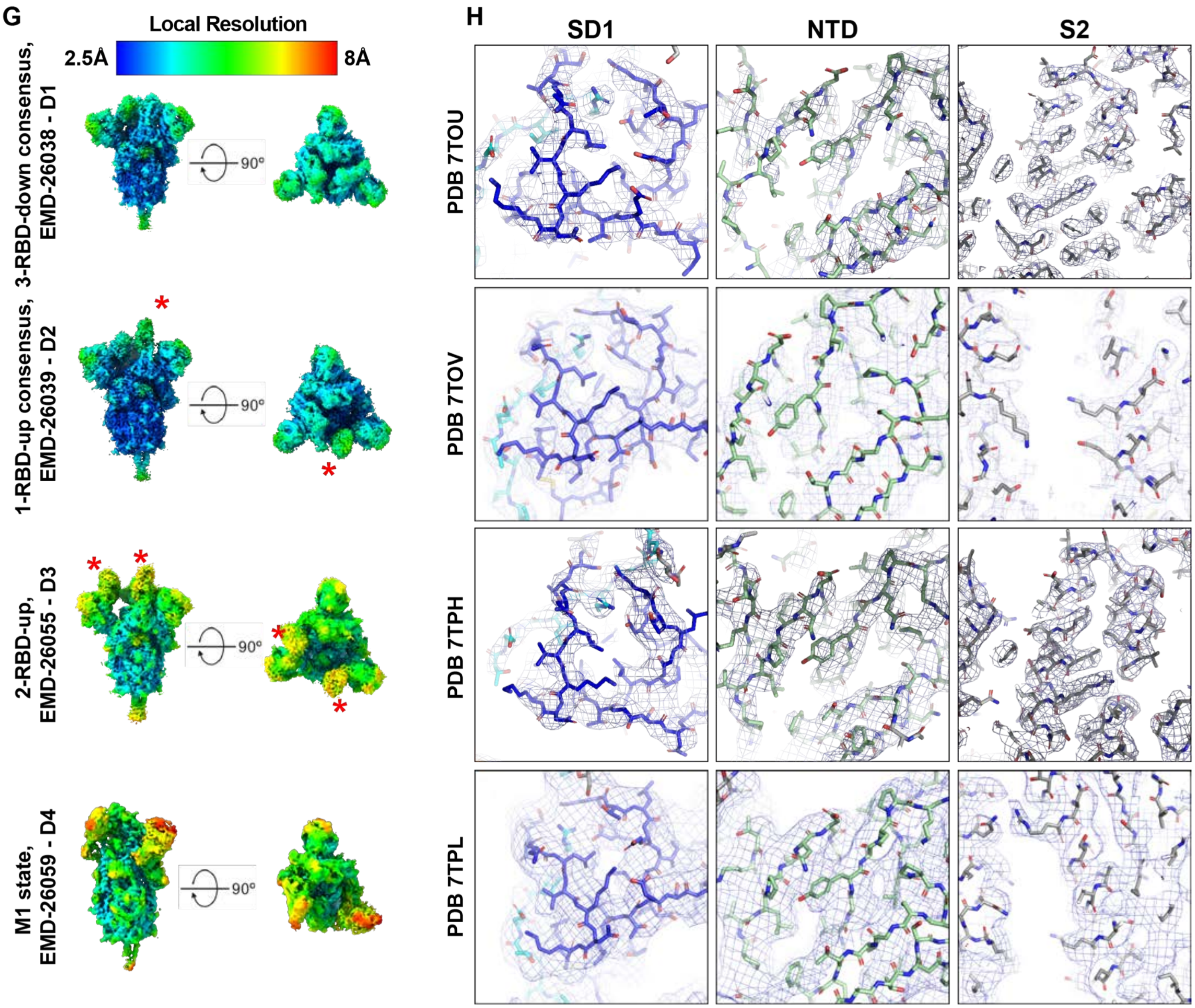


**Fig. S7. Quality assessment of the cryo-EM map and model fitting for the S-GSAS-Delta SARS-CoV-2 S ectodomains. (Left)** Refined maps colored by local resolution ranging from 2.5Å to 8Å**. (Right)** Zoomed in views of SD1, NTD, and S2.

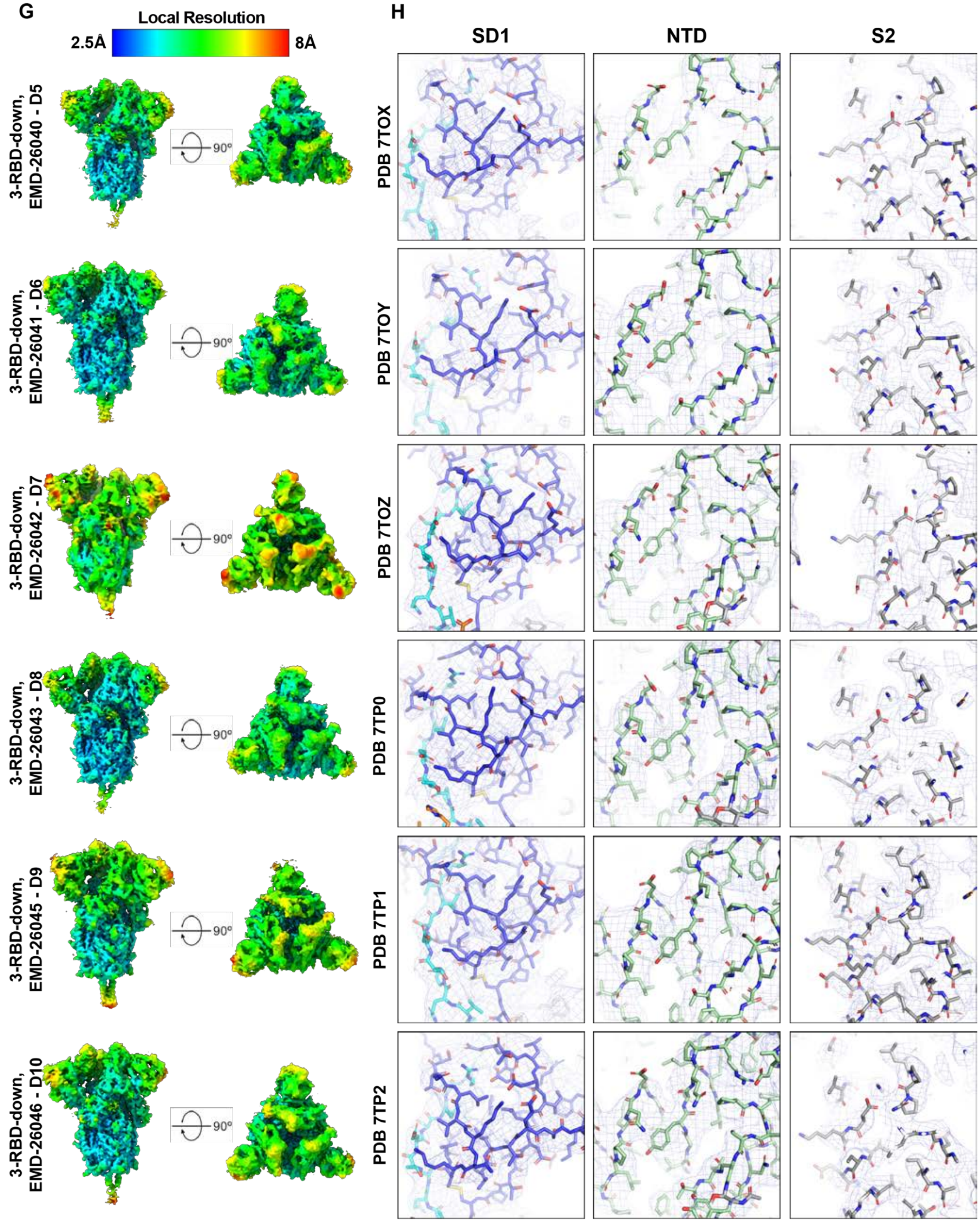


**Fig. S7 (contd). Quality assessment of the cryo-EM map and model fitting for the S-GSAS-Delta SARS-CoV-2 S ectodomains. (Left)** Refined maps colored by local resolution ranging from 2.5Å to 8Å**. (Right)** Zoomed in views of SD1, NTD, and S2.

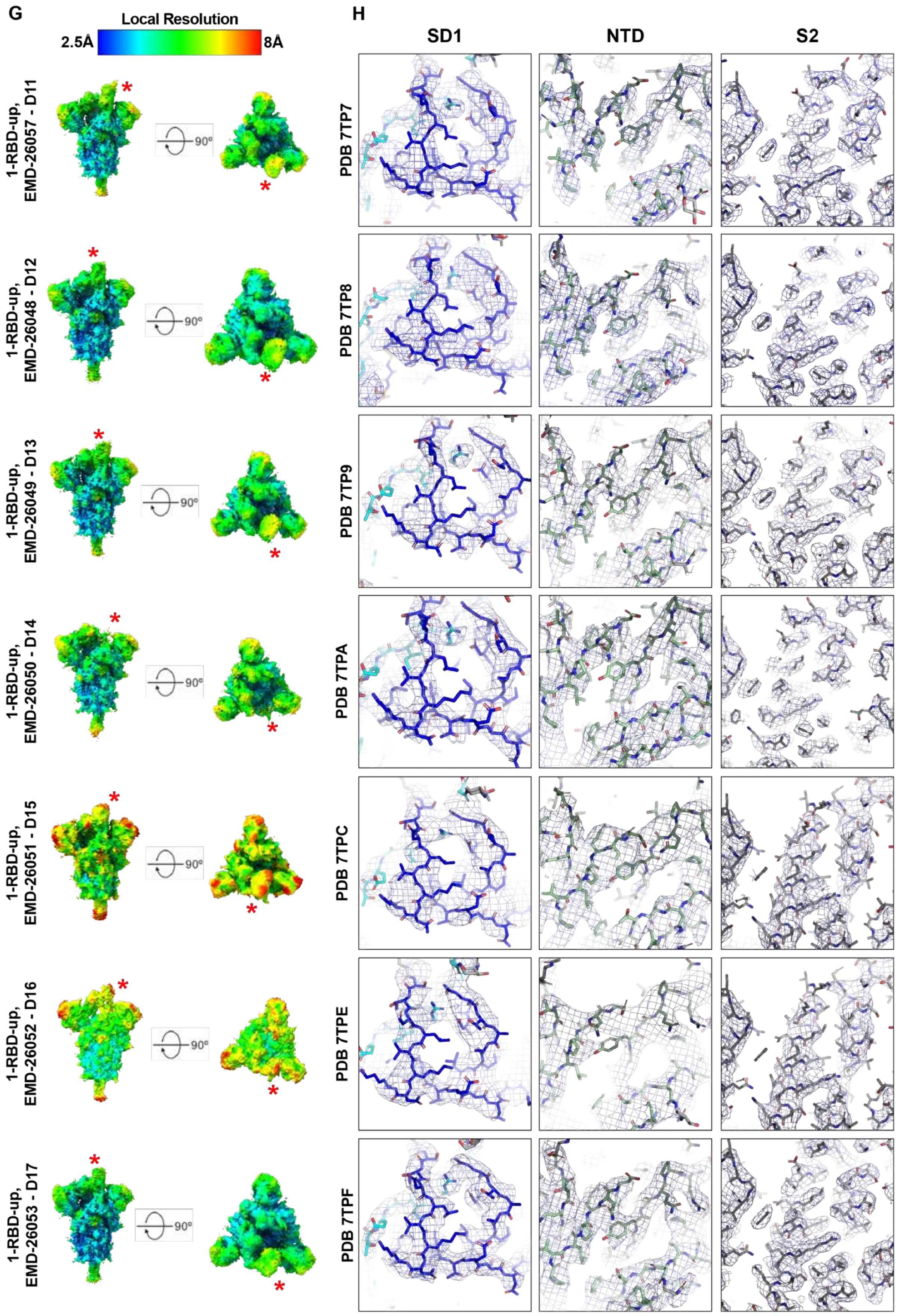


**Fig. S7 (contd.). Quality assessment of the cryo-EM map and model fitting for the S-GSAS-Delta SARS-CoV-2 S ectodomains. (Left)** Refined maps colored by local resolution ranging from 2.5Å to 8Å**. (Right)** Zoomed in views of SD1, NTD, and S2.

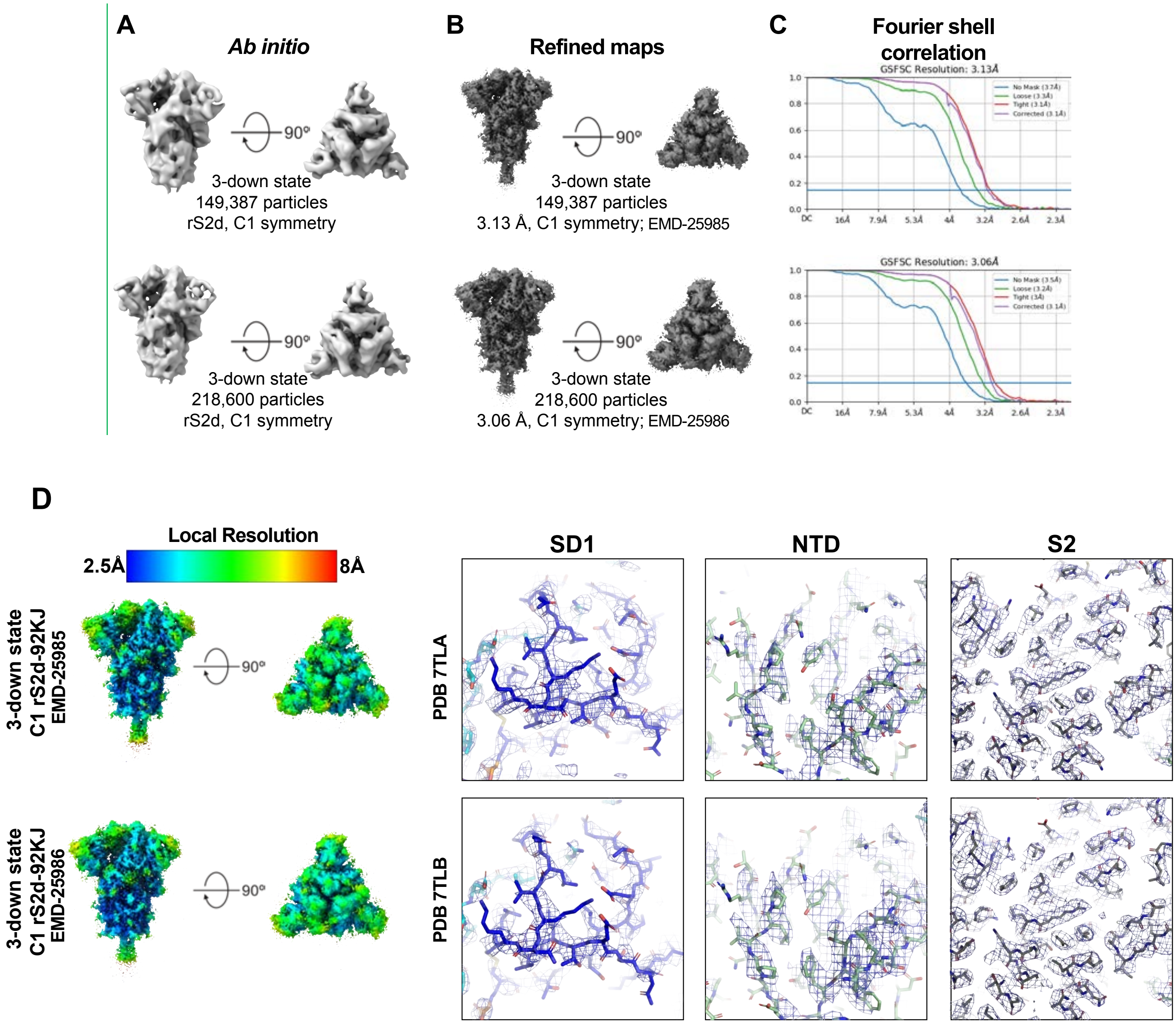


**Fig. S8. Cryo-EM map resolutions for the rS2d SARS-CoV-2 S ectodomains.**
**(A)** *Ab initio* reconstructions for the cryo-EM 3-down states. **(B)** Refined maps for the cryo-EM corresponding states. **(C)** Fourier shell correlation curves for the cryo-EM corresponding states. **(D)** **(Left)** Refined maps colored by local resolution ranging from 2.5Å to 8Å**. (Right)** Zoomed in views of SD1, NTD, and S2.

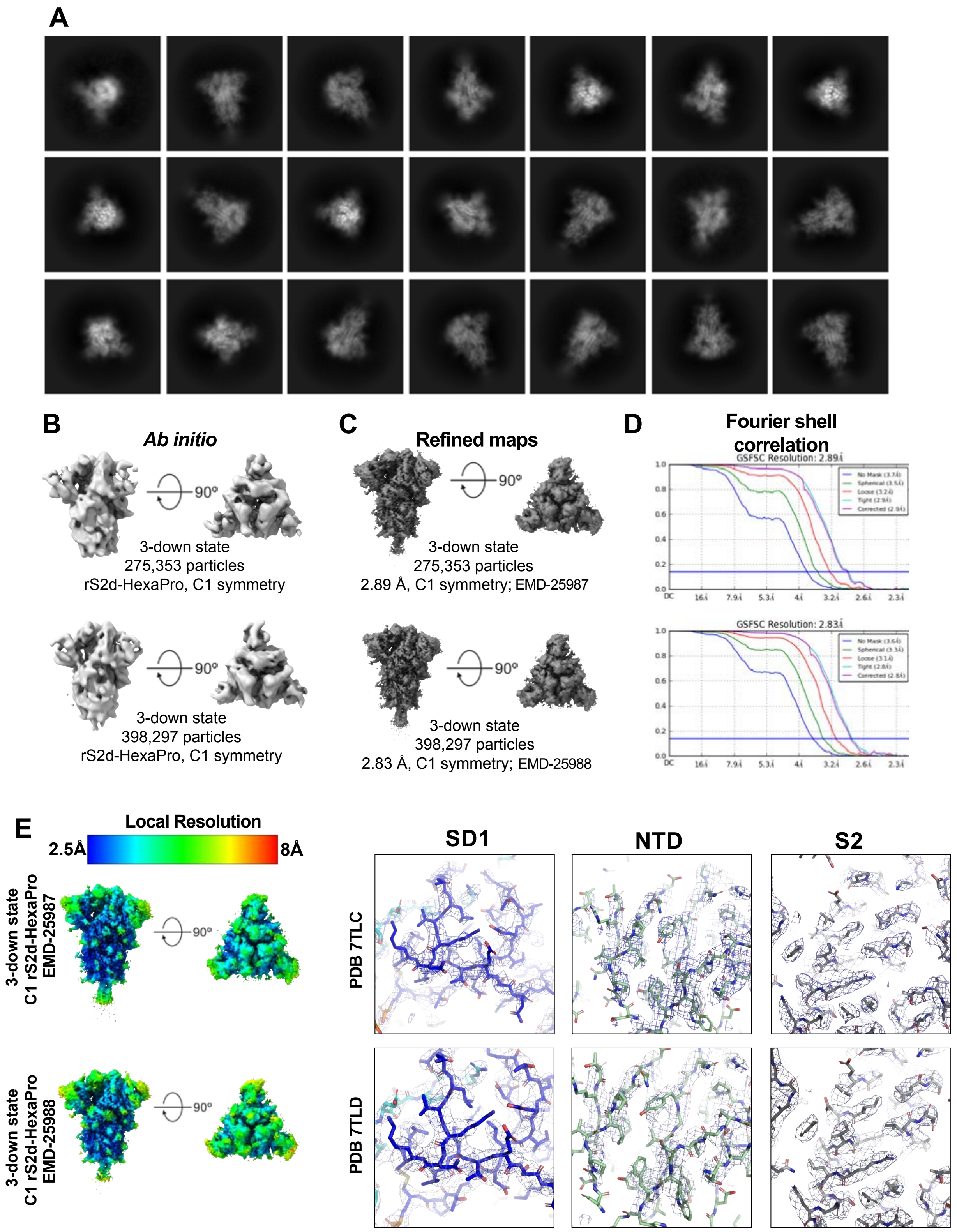


**Fig. S9. 3D reconstructions of the rS2d-HexaPro SARS-CoV-2 S ectodomain.**
**(A)** Representative 2D class averages from Cryo-EM dataset. **(B)** *Ab initio* reconstructions. **(H)** Refined maps for the corresponding cryo-EM states. **(I)** Fourier shell correlation curves for the corresponding cryo-EM states. **(E) (Left)** Refined maps colored by local resolution ranging from 2.5Å to 8Å**. (Right)** Zoomed in views of SD1, NTD, and S2.

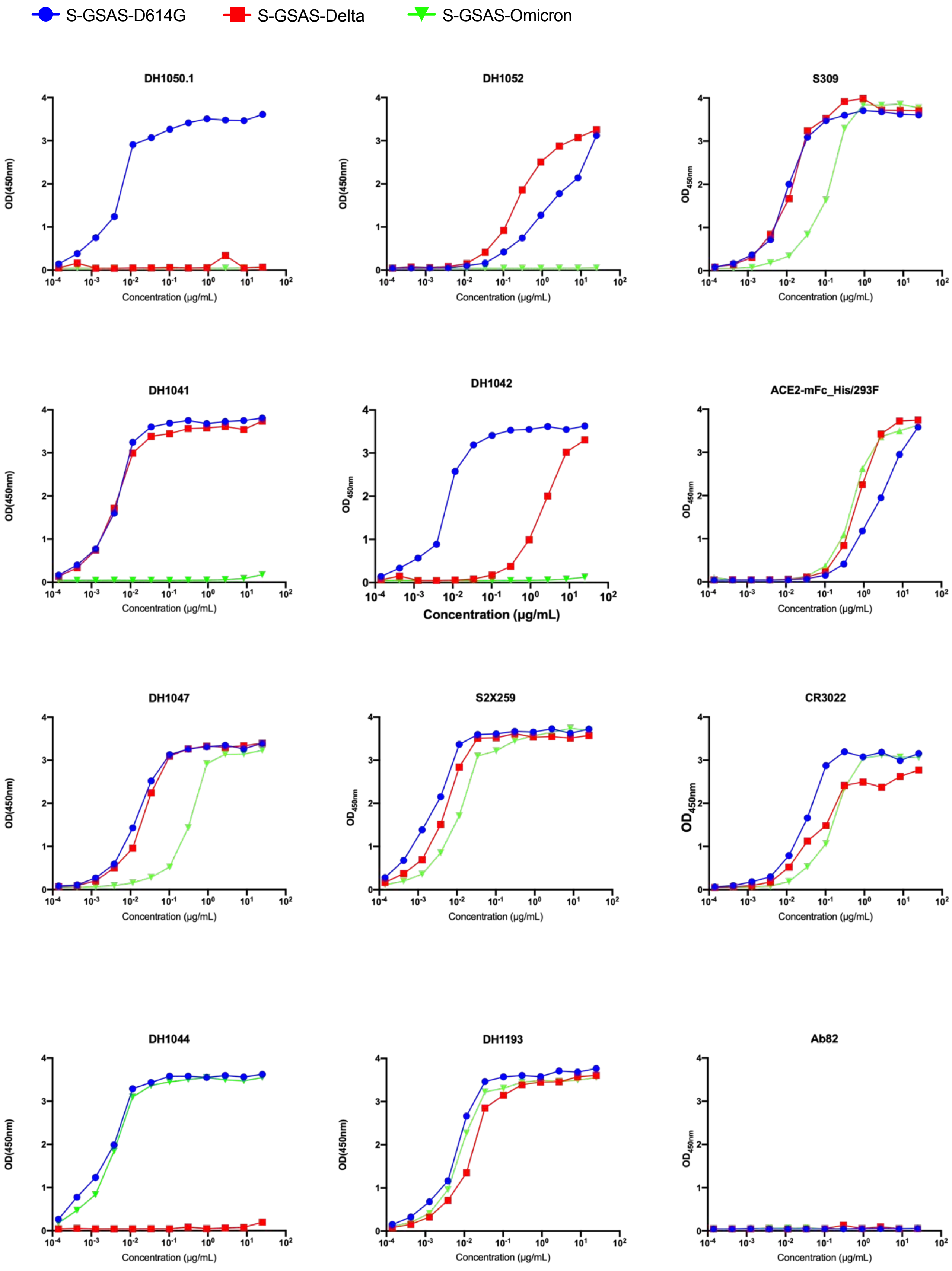

S-GSAS-D614G
S-GSAS-Delta
S-GSAS-Omicron
DH1050.1
DH1052
S309
DH1041
DH1042
ACE2-mFc_His/293F
DH1047
S2X259
CR3022
DH1044
DH1193
Ab82
OD(450nm)
$OD_{450nm}$
Concentration (μg/mL)
0
1
2
3
4
$10^{-4}$
$10^{-3}$
$10^{-2}$
$10^{-1}$
$10^{0}$
$10^{1}$
$10^{2}$

**Fig. S10. Binding of ACE2 receptor ectodomain, and antibodies DH1041, DH1042, DH1044, DH1047, DH1193, S2X259, S309, and CR3022 (RBD-directed), DH1050.1 and DH1052 (NTD-directed) to S protein variants measured by ELISA.** Serially diluted spike protein was bound in individual wells of 384-well plates, which were previously coated with streptavidin. Proteins were incubated and washed, then antibodies at 10µg/ml or ACE2 with a mouse Fc tag at 2µg/ml were added. Antibodies were incubated, washed and binding was detected with goat anti-human-HRP. Data shown are representative of two independent experiments. Ab82 was used as negative control.

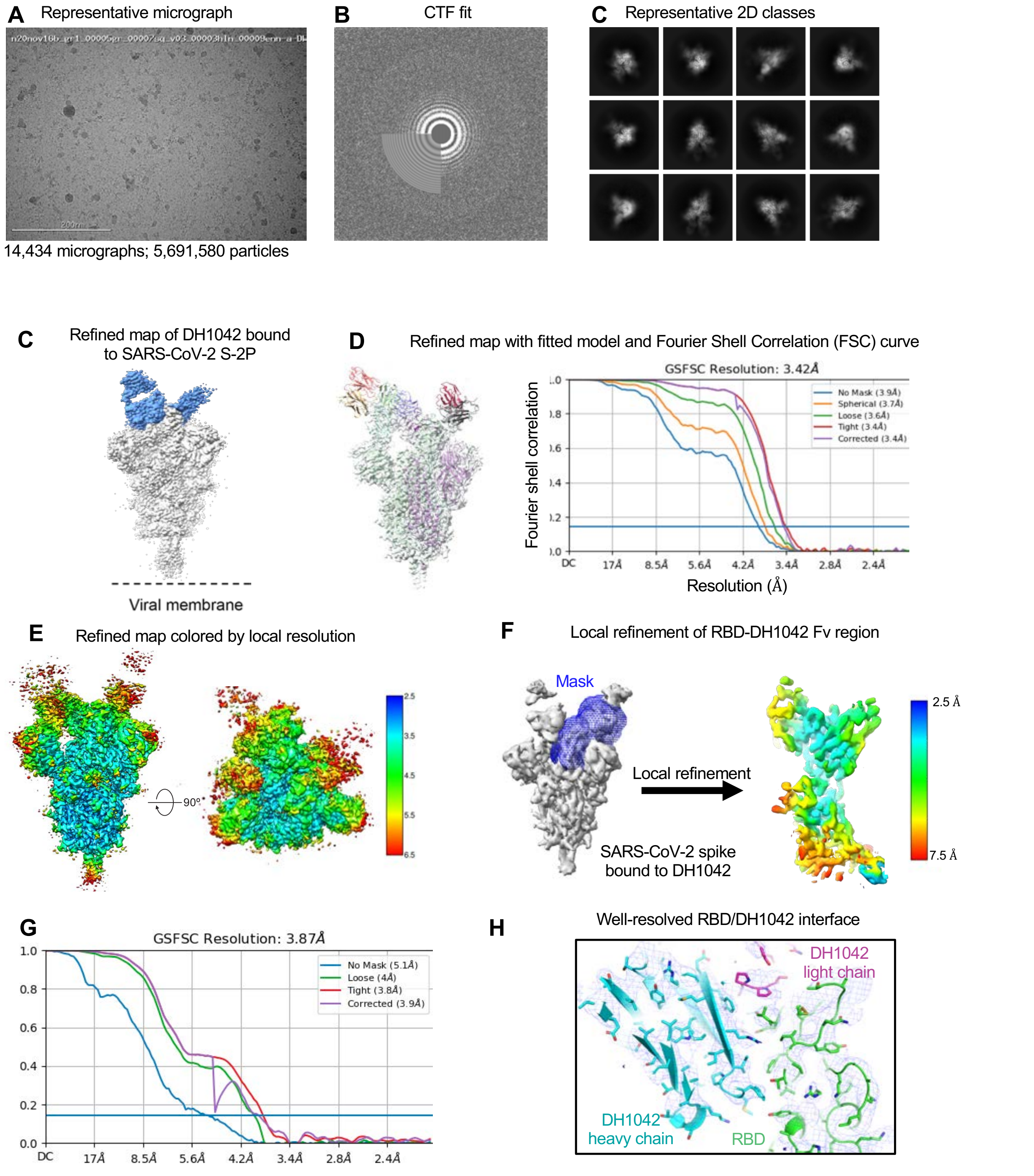


**Figure S11. Cryo-EM data processing for the DH1042 in complex with SARS-CoV-2 S protein.** (**A**) Representative micrograph. (**B**) Representative 2D class averages. (**C**) Refined 3D map segmented and colored by component, with the SARS-CoV-2 S protein colored in grey and DH1042 colored blue. (**D**) Left. Refined map with fitted model. Right. Fourier Shell Correlation (FSC) curves of the 3D reconstructions with horizontal blue line indicating $FSC_{0.143}$. (**E**) Refined 3D density map colored by local resolution in side and top views. (**F**) Local refinement of RBD-DH1042 Fv region. Left Blue mesh shows the mask that was used for local refinement. Right. Local Refined 3D density map colored by local resolution . (**G**) FSC curves of local refined map shown in (F) with horizontal blue line indicating $FSC_{0.143}$. (**H**) View of RBD-DH1042 interface. Cryo-EM reconstruction is shown as blue mesh with underlying fitted model. RBD is colored green, DH1042 heavy chain cyan and DH1042 light chain pink.

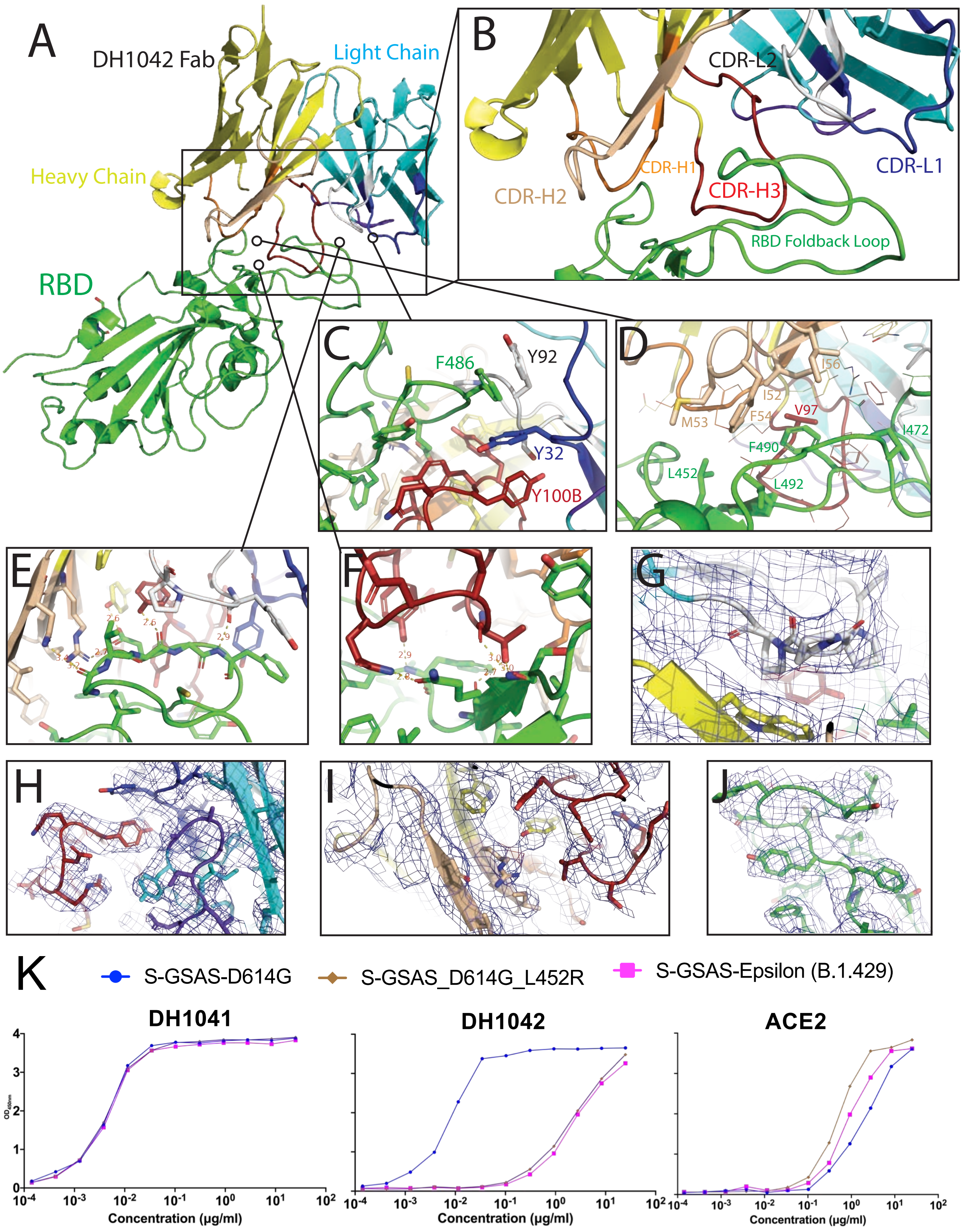
A
DH1042 Fab
Light Chain
Heavy Chain
RBD
B
CDR-L2
CDR-H1
CDR-H2
CDR-H3
CDR-L1
RBD Foldback Loop
C
F486
Y92
Y32
Y100B
D
I56
I52
M53
F54
V97
I472
F490
L452
L492
E
F
G
H
I
J
K
S-GSAS-D614G
S-GSAS_D614G_L452R
S-GSAS-Epsilon (B.1.429)
DH1041
DH1042
ACE2
Concentration (µg/ml)
Concentration (µg/ml)
Concentration (µg/ml)

**Fig. S12. Binding interface of DH1042 Fab and SARS-CoV-2 Spike RBD**. **(A,B)** Cartoon representation of the interface. Green: RBD, Cyan: Fab Light Chain, Yellow: Fab Heavy Chain, Blue: CDR-L1, White: CDR-L2, Purple: CDR-L3 (L3 is not involved in the interface), Orange: CDR-H1, Tan: CDR-H2, Red: CDR-H3. **(C)** A pi-pi stacking interaction comprised of aromatic residues from the RBD, and the L1, L2, and H3 Fab loops. **(D)** A hydrophobic interaction comprised of non-polar residues from the RBD and H2 loop, with additional participation by the H3 loop. Leucine at residue position 452 (L452) is substituted by an Arginine in naturally occurring variants, including Epsilon (B.1.429) and Delta (B.1.617.2). **(E)** Polar interaction centered on the RBD foldback loop. Both Heavy and Light chain of the Fab participate in this interaction and many of the involved RBD atoms are on the main chain. **(F)** Polar interaction at the center of the RBD, closest to the ACE2-binding residues. T98, Q100A, and Y100C of the H3 loop form hydrogen bonds with the RBD, mostly to main chain atoms. **(G-J)** Cryo-EM map shown as mesh surface over fitted model as cartoon representation with key sidechains shown as sticks. **(G)** A triple proline sequence in the L2 loop of the Fab light chain. **(H)** Interface between Fab Light and Heavy chains, highlighting the model's resolution of R99, Y100B (heavy chain), Y32, and Y49 (light chain). **(I)** Fab side chains involved in hydrogen bonding to RBD, including Y100D (heavy chain), R50, and Y95 (light chain). **(J)** Foldback loop of RBD, highlighting residues V483, F486, Y489, and F490. **(K)** Binding of DH1041, DH1042 and ACE2 to S-GSAS-D614G, S-GSAS-L452R, S-GSAS-Epsilon. DH1042 binding is substantially reduced by the L452R substitution in the RBD, and is responsible for its loss in activity against the Epsilon and Delta variants that harbor the L452R substitution.

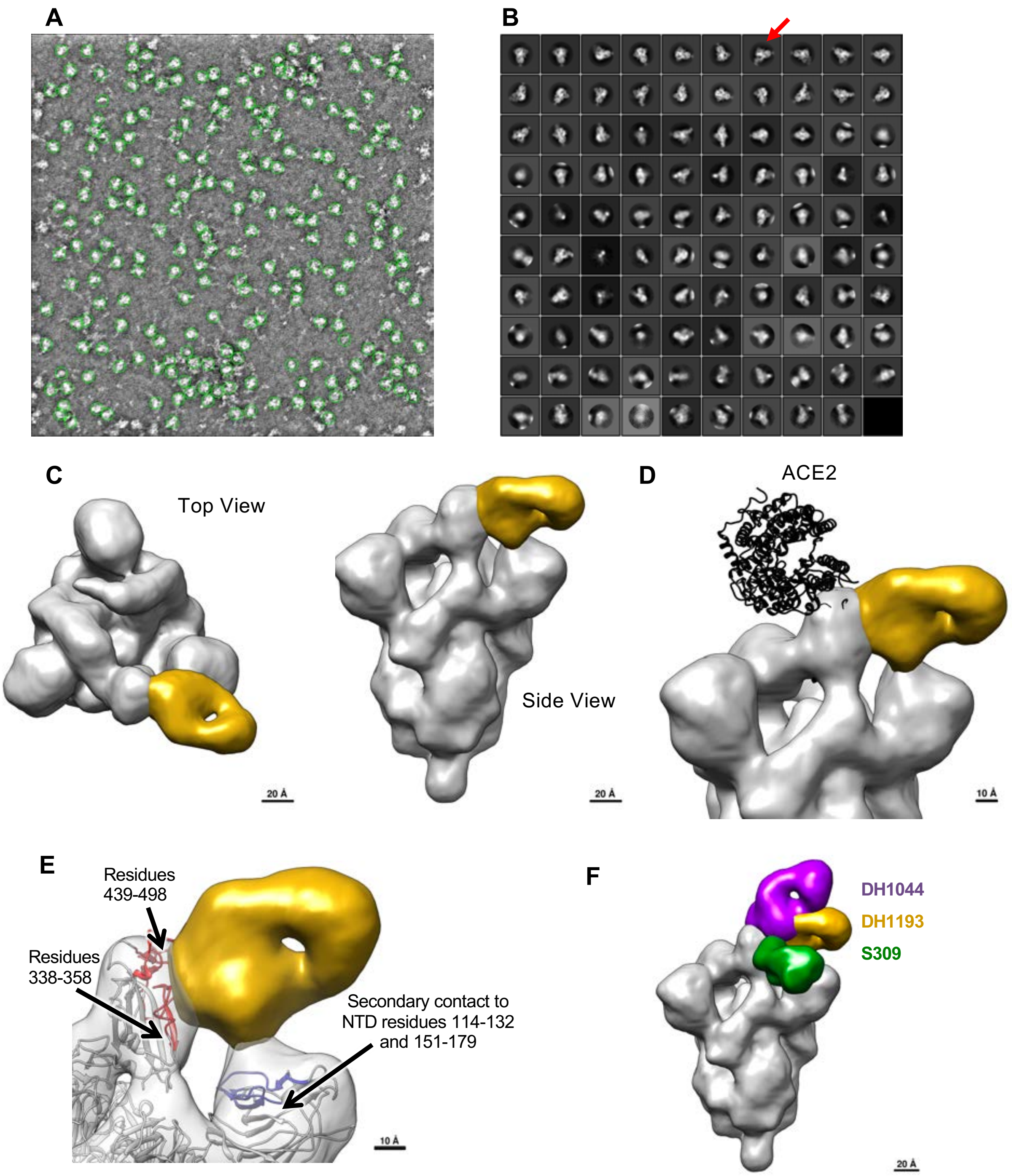


**Fig. S13. NSEM reconstruction of DH1193 Fab bound to SARS-CoV-2 Hexapro spike**. **(A)** Representative micrograph **(B)** Representative 2D classes. Red arrow points to a class showing antibody bound to the spike. **(C)** Left. Top view and Right. Side view of an NSEM reconstruction of SARS-CoV-2 Hexapro spike (gray) bound to DH1193 Fab (yellow). **(D)** Model of DH1193 bound spike overlaid on ACE2 bound spike. ACE2 is shown in black. The modeling indicates that both DH1193 and ACE2 can bind simultaneously to the S protein. **(E)** The DH1193 epitope maps to residues 439-498 and 338-358 of the RBD. Secondary contacts with the adjacent NTD are observed within residues 114-132 and 151-179. **(F)** Overlay showing the locations of S309 (green), DH1193 (mustard) and DH1044 (purple) binding to SARS-CoV-2 S protein RBD.

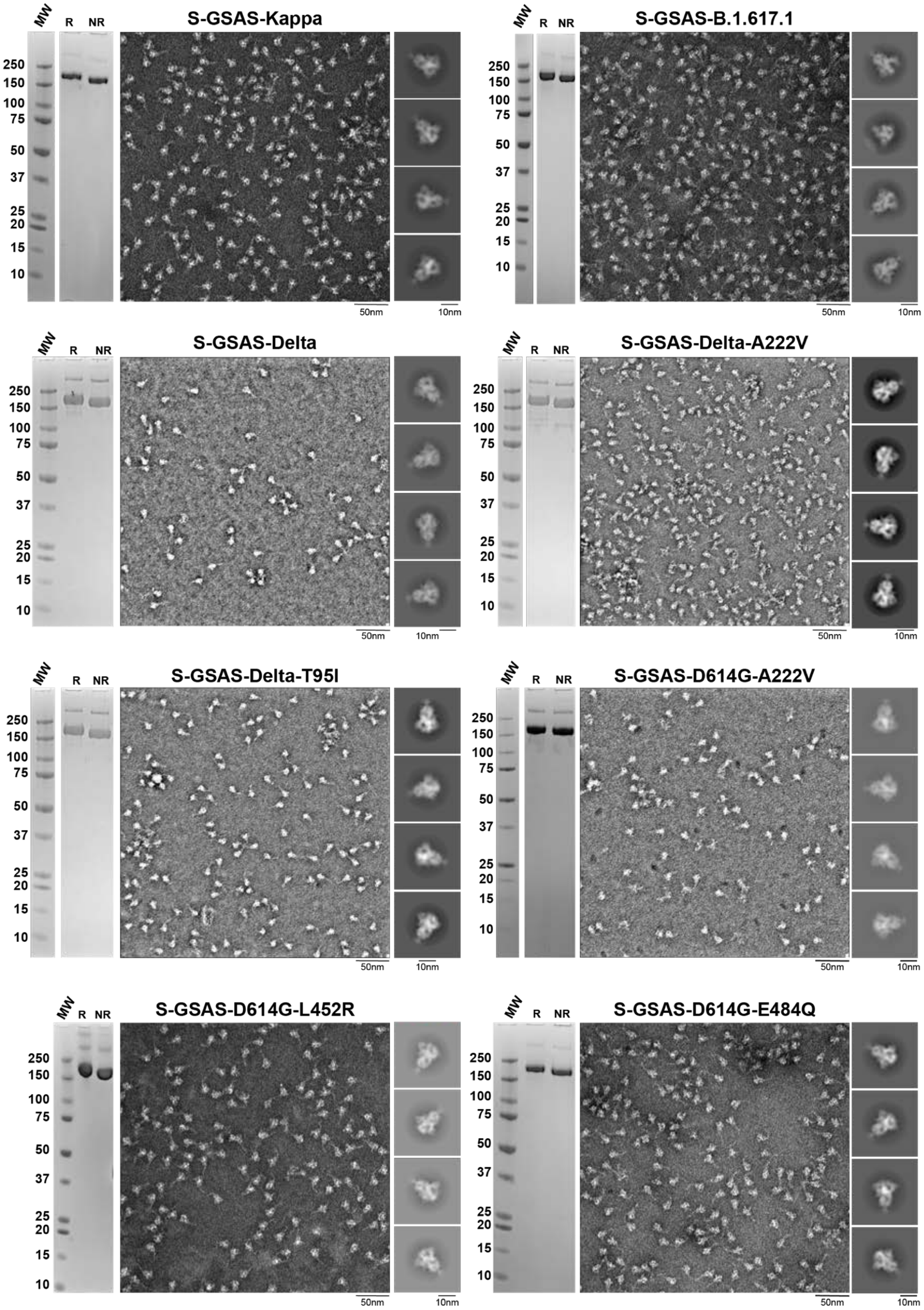


**Fig. S14. Quality control assessment of the SARS-CoV-2 spike variants preparations.** SDS-PAGE of the purified SARS-CoV-2 S protein ectodomains, labels represent: R (Reduced) and NR (Non-Reduced), and representative NSEM micrographs and 2D class averages.

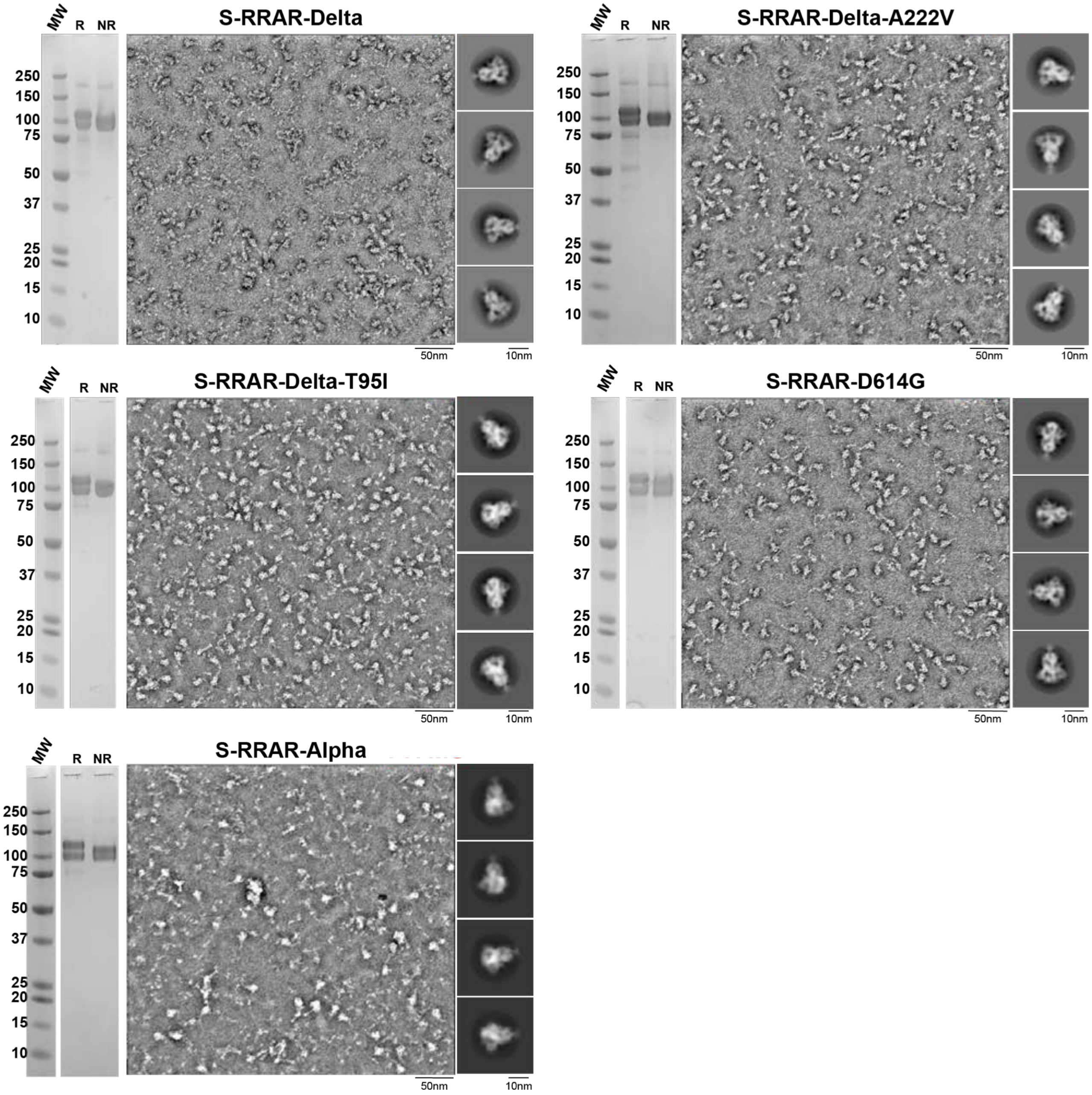


**Fig. S14 (contd.). Quality control assessment of the SARS-CoV-2 spike variants preparations.** SDS-PAGE of the purified SARS-CoV-2 S protein ectodomains, labels represent: R (Reduced) and NR (Non-Reduced), and representative NSEM micrographs and 2D class averages.

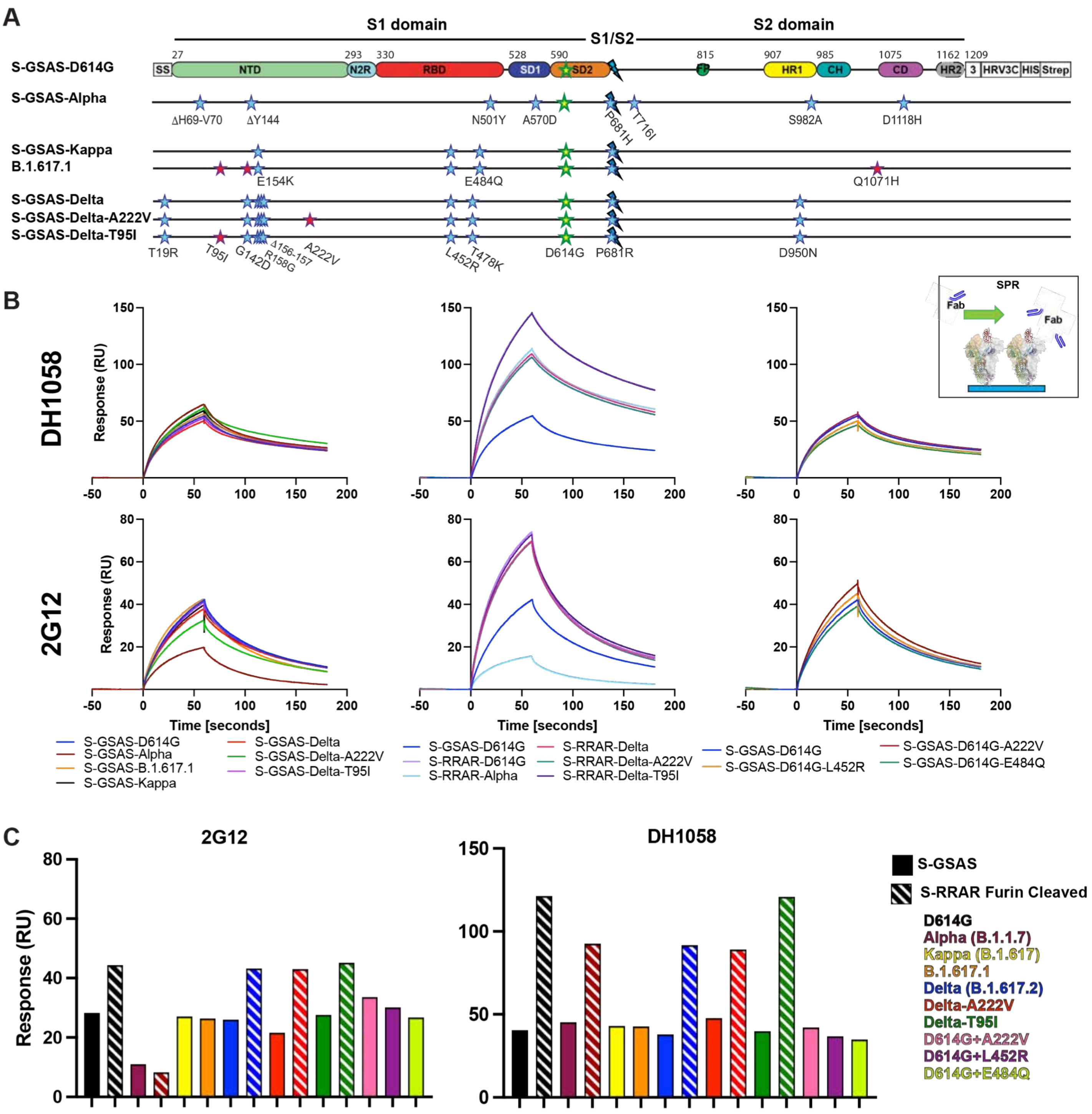
A
S1 domain
S1/S2
S2 domain
S-GSAS-D614G
S-GSAS-Alpha
S-GSAS-Kappa
B.1.617.1
S-GSAS-Delta
S-GSAS-Delta-A222V
S-GSAS-Delta-T95I
SS NTD N2R RBD SD1 SD2 FP HR1 CH CD HR2 3 HRV3C HIS Strep
27 293 330 528 590 815 907 985 1075 1162 1209
ΔH69-V70 ΔY144 N501Y A570D P681H T716I S982A D1118H
E154K E484Q Q1071H
T19R T95I G142D Δ156-157 R158G A222V L452R T478K D614G P681R D950N
B
DH1058
2G12
Response (RU)
Time [seconds]
SPR
Fab
S-GSAS-D614G
S-GSAS-Alpha
S-GSAS-B.1.617.1
S-GSAS-Kappa
S-GSAS-Delta
S-GSAS-Delta-A222V
S-GSAS-Delta-T95I
S-RRAR-D614G
S-RRAR-Alpha
S-RRAR-Delta
S-RRAR-Delta-A222V
S-RRAR-Delta-T95I
S-GSAS-D614G-L452R
S-GSAS-D614G-A222V
S-GSAS-D614G-E484Q
C
2G12
DH1058
Response (RU)
S-GSAS
S-RRAR Furin Cleaved
D614G
Alpha (B.1.1.7)
Kappa (B.1.617)
B.1.617.1
Delta (B.1.617.2)
Delta-A222V
Delta-T95I
D614G+A222V
D614G+L452R
D614G+E484Q

**Figure S15. Binding of DH1058, an FP-directed antibody, and 2G12, an S2-glycan directed antibody, to spike variants measured by SPR. (A)** Substitution of the SARS-CoV-2 variants. The subunits are identified as in Figure 1. The D614G mutation is in the SD2 domain (yellow star, green contour). The S1/S2 furin cleavage site (RRAR) has either been maintain (S-RRAR) or been mutated to GSAS (S-GSAS). The substitutions in each variant are indicated by blue (shared) and red (differing within a set *i.e*. Kappa/B.1.617.1 and Delta) stars. Multiple ectodomain constructs in the S-GSAS-Delta spike backbone were tested that differed in their NTD mutations (A222V and T95I). **(B)** Binding of antibodies DH1058 and 2G12, to spike variants measured by SPR. The schematic shows the assay format. For the SPR, spike variants were captured on a Series S Strepavidin (SA) chip coated at 200 nM. Fabs were then injected at 200nM with a contact time of 60s and a dissociation time of 120s (50µL/min). The S-RRAR S ectodomain where cotransfected with furin to cleave the S protein at the S1/S2 junction during expression (see Figure S14 for QC analysis). **(C)** The bar graphs represent the response levels (RU) after 20 seconds of dissociation.

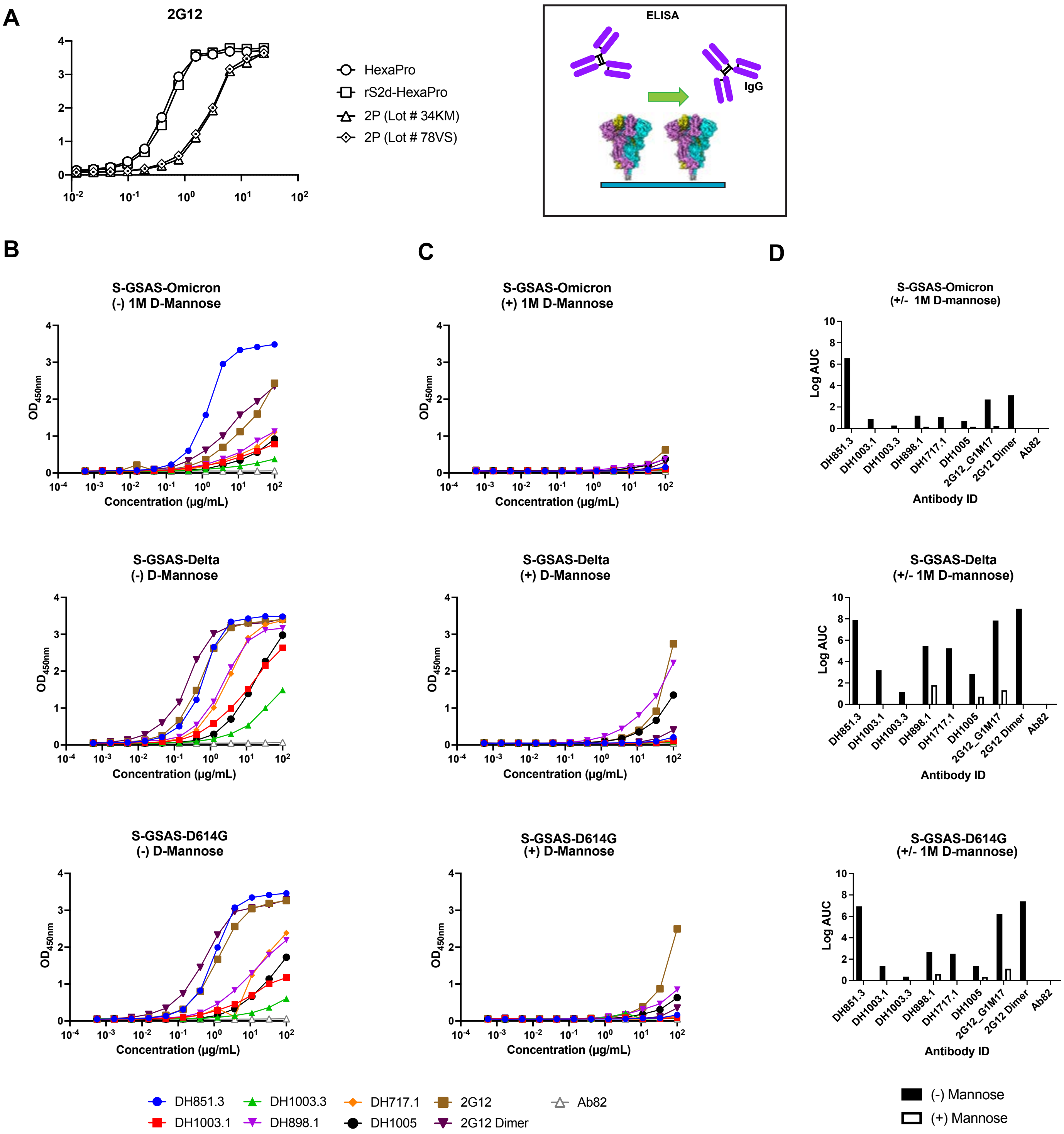

A
2G12
HexaPro
rS2d-HexaPro
2P (Lot # 34KM)
2P (Lot # 78VS)
ELISA
IgG
B
C
D
S-GSAS-Omicron (-) 1M D-Mannose
S-GSAS-Omicron (+) 1M D-Mannose
S-GSAS-Omicron (+/- 1M D-mannose)
S-GSAS-Delta (-) D-Mannose
S-GSAS-Delta (+) D-Mannose
S-GSAS-Delta (+/- 1M D-mannose)
S-GSAS-D614G (-) D-Mannose
S-GSAS-D614G (+) D-Mannose
S-GSAS-D614G (+/- 1M D-mannose)
$OD_{450nm}$
Concentration (μg/mL)
Log AUC
Antibody ID
DH851.3
DH1003.1
DH1003.3
DH898.1
DH717.1
DH1005
2G12_G1M17
2G12 Dimer
Ab82
2G12
(-) Mannose
(+) Mannose

**Figure S16. Binding of Fab-dimerized glycan-reactive (FDG) antibodies to SARS-CoV-2 S ectodomains. A.** Binding of 2G12 to SARS-CoV-2 S-2P and Hexapro. Schematic shows assay format. **B-C.** Binding of FDG antibodies to SARS-CoV-2 S ectodomain either **B.** without or **C.** in presence of D-mannose. **D**. The bar graphs represent area under the curve calculated from ELISA plots shown in panels **B** and **C**. Black and white bars show binding levels without and with D-mannose, respectively.

**Table S2 Crystallographic Data Collection and Refinement Statistics for DH1058 Fab bound to SARS-CoV-2 fusion peptide**

| PDB Accession Code | 7TOW |
|---|---|
| Space group | *P 1 21 1* |
| Unit-cell dimensions | |
| a, b, c (Å) | 53.9, 76.8, 119.8 |
| α, β, $\Upsilon$ (º) | 90.0, 100.9, 90.0 |
| Resolution (Å) | 2.15 (2.15-38.96) |
| CC(1/2) | 0.960 (0.993) |
| CC* | 0.990 (0.998) |
| Rpim | 0.115 (0.035) |
| R-work[b] | 0.169 (0.167) |
| R-free[b] | 0.253 (0.234) |
| Overall R-sym | 0.075 |
| I/σ(I) | 5.4 (22.5) |
| Completeness (%) | 95.1 (92.8) |
| No. reflection/unique | 4,817 (48,437) |
| Refinement | |
| Molprobity | |
| Ramachandran favored | 95.68 |
| Ramachandran allowed | 3.24 |
| Ramachandran outlier | 1.08 |
| Rotamer outliers | 4.44 |
| Clash score | 4.35 |
| No. of water | 793 |
| RMSD | |
| Bond lengths (Å) | 0.007 |
| Bond angles (°) | 0.94 |

Values are listed for highest resolution shell. Values in parentheses are overall statistics. $R_{free}$ is calculated from 5% of the reflections excluded from refinement.
[a]$R_{pim}$ = Sum(h) [ Sqrt(1/(n-1)) Sum(j) [I(hj) - <Ih>] / Sum(hj) <Ih>
where n is the number of observations of reflection h (i.e., j=1,n)
[b]R = Σhkl||Fobs| − |Fcalc||/Σhkl|Fobs|.
[a]$R_{sym}$ = Σ|I − ⟨I⟩ |/Σ ⟨I⟩, where I is the observed intensity, and ⟨I⟩ is the average intensity of multiple observations of symmetry-related reflections.
